# Gene duplication and rate variation in the evolution of plastid ACCase and Clp genes in angiosperms

**DOI:** 10.1101/2021.09.13.460099

**Authors:** Alissa M. Williams, Olivia G. Carter, Evan S. Forsythe, Hannah K. Mendoza, Daniel B. Sloan

## Abstract

While the chloroplast (plastid) is known for its role in photosynthesis, it is also involved in many other metabolic pathways essential for plant survival. As such, plastids contain an extensive suite of enzymes required for non-photosynthetic processes. The evolution of the associated genes has been especially dynamic in flowering plants (angiosperms), including examples of gene duplication and extensive rate variation. We examined the role of ongoing gene duplication in two key plastid enzymes, the acetyl-CoA carboxylase (ACCase) and the caseinolytic protease (Clp), responsible for fatty acid biosynthesis and protein turnover, respectively. In plants, there are two ACCase complexes—a homomeric version present in the cytosol and a heteromeric version present in the plastid. Duplications of the nuclear-encoded homomeric ACCase gene and retargeting of one resultant protein to the plastid have been previously reported in multiple species. We find that these retargeted homomeric ACCase proteins exhibit elevated rates of sequence evolution, consistent with neofunctionalization and/or relaxation of selection. The plastid Clp complex catalytic core is composed of nine paralogous proteins that arose via ancient gene duplication in the cyanobacterial/plastid lineage. We show that further gene duplication occurred more recently in the nuclear-encoded core subunits of this complex, yielding additional paralogs in many species of angiosperms. Moreover, in six of eight cases, subunits that have undergone recent duplication display increased rates of sequence evolution relative to those that have remained single copy. We also compared substitution patterns between pairs of Clp core paralogs to gain insight into post-duplication evolutionary routes. These results show that gene duplication and rate variation continue to shape the plastid proteome.

## Introduction

The plastid is a dynamic proteomic environment in which key photosynthetic and non-photosynthetic biochemical reactions occur. Major non-photosynthetic functions of plastids include the reaction catalyzed by the acetyl-CoA carboxylase (ACCase) enzyme and protein degradation performed by the caseinolytic protease (Clp) complex (Caroca et al., 2021; Green, 2011; Konishi et al., 1996; Nishimura et al., 2017; Nishimura and van Wijk, 2015). Both of these functions are essential in plants and thus the genes involved are generally highly conserved; however, these genes have undergone rapid evolution in multiple angiosperm species (Barnard-Kubow et al., 2014; Erixon and Oxelman, 2008; Jansen et al., 2007; Park et al., 2017; Sloan et al., 2014, 2014; Wicke et al., 2011; Williams et al., 2019, 2015; Zhang et al., 2014). While many hypotheses about these patterns of accelerated evolution have been posited, the underlying evolutionary mechanisms, causes, and consequences remain largely unknown.

The ACCase enzyme catalyzes the first committed step of fatty acid biosynthesis, the carboxylation of acetyl-CoA to malonyl-CoA (Salie and Thelen, 2016; Sasaki and Nagano, 2004). This step requires four different enzyme domains—one biotin carboxylase, one biotin carboxyl carrier, and two (α andβ) carboxyltransferases (Salie and Thelen, 2016; Sasaki and Nagano, 2004; Schulte et al., 1997). In most plants, there are two forms of the ACCase enzyme. The homomeric version, present in the cytosol, is encoded by a single nuclear gene (Konishi et al., 1996; Konishi and Sasaki, 1994; Sudianto and Chaw, 2019). The heteromeric version, present in the plastid, is encoded by five genes in *Arabidopsis thaliana*; each functional domain is represented by a single gene except for the biotin carboxyl carrier domain, which is encoded by two genes (Konishi et al., 1996; Konishi and Sasaki, 1994; Salie and Thelen, 2016). Four of these genes are in the nuclear genome while the fifth (*accD*) is in the plastid genome (Caroca et al., 2021; Sasaki and Nagano, 2004). In a few angiosperm lineages, including the Brassicaceae, Caryophyllaceae, Geraniaceae, and Poaceae, there have been duplications of the homomeric ACCase gene with subsequent retargeting of one resultant protein to the plastid (Babiychuk et al., 2011; Konishi and Sasaki, 1994; Park et al., 2017; Parker et al., 2014; Rockenbach et al., 2016; Schulte et al., 1997).

The Clp complex is one of the most abundant stromal proteases and degrades a variety of targets (Apitz et al., 2016; Bouchnak and van Wijk, 2021; Majeran et al., 2000; Montandon et al., 2019; Nishimura et al., 2017; Nishimura and van Wijk, 2015; Welsch et al., 2018). This complex consists of many types of subunits. Adapters bind proteins targeted for degradation and deliver them to chaperones, which use ATP to unfold the targeted proteins into the proteolytic core of the complex (Nishimura and van Wijk, 2015). The core consists of 14 subunits that are encoded by nine different paralogous genes (Olinares et al., 2011a; Peltier et al., 2004; Sjögren et al., 2006; Stanne et al., 2007). Eight of these genes reside in the nuclear genome (*CLPP3-6, CLPR1-4*), while the ninth is encoded in the plastid genome (*clpP1*) (Nishimura et al., 2017; Olinares et al., 2011b). The ClpP subunits contain a catalytically active Ser-His-Asp triad, whereas the ClpR subunits do not (Nishimura and van Wijk, 2015; Porankiewicz et al., 1999). These nine paralogs are the results of gene duplications throughout cyanobacterial and plastid evolution and are shared by all land plants (Olinares et al., 2011a). Ongoing gene duplication of individual subunits has been noted in a handful of angiosperm lineages (Rockenbach et al., 2016; Williams et al., 2021, 2019).

Thus, the evolutionary trajectory of both of these essential plastid pathways is characterized by gene duplication at both ancient and recent timescales. Gene duplication is common in land plants, in part due to the frequency with which whole genome duplication (polyploidization) occurs in this lineage (Clark and Donoghue, 2018; De Bodt et al., 2005; del Pozo and Ramirez-Parra, 2015; Flagel and Wendel, 2009; Panchy et al., 2016; Wendel et al., 2018). Nearly all species of land plants have polyploidization events in their evolutionary histories (Clark and Donoghue, 2018; Leebens-Mack et al., 2019; Panchy et al., 2016). Angiosperms in particular seem to have a propensity for whole genome duplication; the entire clade shares an ancient polyploidization event and many lineages have undergone subsequent rounds of whole genome duplication (Clark and Donoghue, 2018; Panchy et al., 2016; Renny-Byfield and Wendel, 2014; Soltis et al., 2009). While every gene is initially affected by whole genome duplication, only 10-30% of those duplicates are maintained in the genome longer-term (Hahn, 2009; Maere et al., 2005; Paterson et al., 2006). Though polyploidy is likely a main contributor to gene duplication in plants, other forms of gene duplication are also prevalent (Flagel and Wendel, 2009). For instance, tandem duplication has been shown to be common in both *Arabidopsis thaliana* and *Oryza sativa*, where tandemly arrayed gene clusters make up 15-20% of genic content. Additionally, multiple studies have shown that transposon-mediated gene duplication occurs frequently in plants (Flagel and Wendel, 2009; Freeling et al., 2008; Rizzon et al., 2006; Wang et al., 2006).

Gene duplication is an important evolutionary process and is thought to be a major source of evolutionary novelty (Hahn, 2009; Ohno, 1970; Taylor and Raes, 2004; Zhang, 2003). The most common evolutionary fate of paralogs is retention of one copy and pseudogenization and loss of the other copy (Lynch and Conery, 2000; Zhang, 2003; Zhang et al., 2003). However, several evolutionary mechanisms have been described in which retention of both gene duplicates is favored. The increased gene-dosage advantage model describes a scenario in which increased amount of gene product produced by the two essentially identical gene copies is beneficial and thus both copies retain ancestral function (Hahn, 2009; Ohno, 1970; Pegueroles et al., 2013; Zhang, 2003). The neofunctionalization model posits that one paralog acquires new functions while the other retains ancestral functionality (Hahn, 2009; Ohno, 1970; Pegueroles et al., 2013; Zhang, 2003). In the subfunctionalization model, an ancestral function is split between the two duplicates (Hahn, 2009; Ohno, 1970; Pegueroles et al., 2013; Zhang, 2003), in some cases creating the possibility for each paralog to optimize a subset of the ancestral function in a process known as escape from adaptive conflict (Des Marais and Rausher, 2008; Huang et al., 2015; Sikosek et al., 2012).

To distinguish between these evolutionary fates, many studies have employed evolutionary rate comparisons (Hahn, 2009; Pegueroles et al., 2013). These comparisons involve both paralogs as well as their common ancestor (Pegueroles et al., 2013). Under both the gene-dosage advantage and subfunctionalization models, gene duplicates are expected to evolve at approximately the same rate as each other (Pegueroles et al., 2013). The difference in evolutionary rates predicted by these two models is found in comparisons to the common ancestor; with a gene-dosage advantage, the expectation is that the paralogs will evolve at the same rate as the common ancestor, while with subfunctionalization, the expectation is that the paralogs will evolve at an increased rate relative to the common ancestor (though this assumption has been challenged) (Force et al., 1999; Hahn, 2009; He and Zhang, 2005; Lynch and Force, 2000; Pegueroles et al., 2013; Zhang, 2003). By contrast, under the neofunctionalization model, asymmetry between evolutionary rates of paralogs is expected, where one paralog retains the ancestral evolutionary rate while the other experiences rate acceleration after being freed from selective constraints (Hahn, 2009; Pegueroles et al., 2013; Zhang, 2003). The proportion of paralogs with asymmetric rates of evolution has been estimated at anywhere from 5% to 65% in a variety of studies (Conant and Wagner, 2003; Dermitzakis and Clark, 2001; Kondrashov et al., 2002; Panchin et al., 2010; Pegueroles et al., 2013; Van de Peer et al., 2001). This wide range of estimates is likely due to differences in study systems, definitions and identifications of paralogs, gene types, and time since duplication. Despite the varying estimates of evolutionary rate asymmetry, it is clear that paralogs evolve under a mixture of evolutionary regimes.

In the case of duplications of the homomeric ACCase gene (*ACC*) and subsequent retargeting of one protein to the plastid, the process of protein retargeting is inherently a form of neofunctionalization because the newly retargeted protein functions in a different cellular compartment than it did ancestrally. As outlined above, a hallmark of neofunctionalization is an increased evolutionary rate in the neofunctionalized gene copy (Hahn, 2009; Pegueroles et al., 2013; Zhang, 2003). Notably, however, relaxed selection and/or pseudogenization can also produce a signature of evolutionary rate asymmetry between paralogs (Lynch and Conery, 2000; Zhang, 2003; Zhang et al., 2003).

The retention of plastid Clp core subunit duplicates is likely due, at least in part, to subfunctionalization. The Clp complex is widely conserved across bacteria; in most bacteria, including *E. coli*, the Clp core consists of 14 identical subunits (Nishimura and van Wijk, 2015; Yu and Houry, 2007). However, in cyanobacteria, several duplications have produced four core subunits—three catalytic ClpP subunits and one catalytically inactive ClpR subunit (Andersson et al., 2009; Stanne et al., 2007). In green lineage (Viridiplantae) plastids, which are descended from ancient cyanobacteria, gene duplication has continued to expand the number of core subunit genes to nine, as described above (Nishimura and van Wijk, 2015). Interestingly, ClpR subunits are incorporated into the core despite their lack of catalytic activity; they are thought to play a structural role in the complex, including chaperone docking (Nishimura and van Wijk, 2015; Olinares et al., 2011b, 2011a; Sjögren and Clarke, 2011). In *A. thaliana*, the Clp chaperone is believed to bind only to the ClpP1/ClpR1-4 ring, whereas chaperone proteins bind to both rings of the Clp core in bacteria (Peltier et al., 2004; Yu and Houry, 2007). This ClpP/ClpR division of function (catalytic activity vs chaperone binding) is indicative of subfunctionalization. Further, though the plastid Clp core subunit genes share common ancestry and are structurally similar, knockouts of individual subunits tend to produce severe phenotypes, including lethality in several cases (Kim et al., 2009; Koussevitzky et al., 2007; Rudella et al., 2006)

Here, we characterize recent gene duplication events and subsequent changes in evolutionary rate in ACCase and Clp core subunits. We show that ACCase genes exhibit patterns of duplication, protein retargeting, and accelerated evolution consistent with neofunctionalization and/or relaxed selection. Additionally, we examine duplications of nuclear-encoded plastid Clp core subunits and demonstrate that duplication leads to significant changes in the rate of evolution in most cases but that patterns differ across Clp subunits, meaning multiple post-duplication evolutionary routes are represented across pairs of paralogs. This work provides additional insights into the interplay between gene duplication and evolutionary rate in the molecular evolution of plastid proteins.

## Materials and Methods

### Compilation and curation of *ACC* nucleotide sequences

Previous work identified duplications of the homomeric ACCase gene *ACC* and subsequent retargeting of one resultant protein to the plastid in the angiosperm families Poaceae (Konishi and Sasaki, 1994; Park et al., 2017; Rockenbach et al., 2016), Brassicaceae (Babiychuk et al., 2011; Park et al., 2017; Parker et al., 2014; Schulte et al., 1997), Caryophyllaceae (Rockenbach et al., 2016), and Geraniaceae (Park et al., 2017). *ACC* sequences were obtained for multiple species in each of these families. All *ACC* genes encoding cytosol-targeted proteins were designated *ACC1* while all *ACC* genes encoding plastid-targeted proteins were designated *ACC2* per established conventions (Babiychuk et al., 2011; Sasaki and Nagano, 2004); thus, sharing the same identifier does not necessarily indicate orthology because of the multiple independent origins of plastid-targeted *ACC2* genes. *Amborella trichopoda*, which has a single *ACC* gene that we designated *ACC1*, was used as an outgroup.

Trimmed *ACC1* and *ACC2* coding sequences (CDSs) were obtained from Rockenbach et al. (2016) for *Amborella trichopoda, Arabidopsis thaliana, Agrostemma githago*, *Silene noctiflora*, *Silene paradoxa,* and *Triticum aestivum*. The trimming in Rockenbach et al (2016) was codon-guided and included removal of the target peptide. *ACC1* and *ACC2* CDSs from the following species were compiled using gene identifiers from Table S4 in Park et al. (2017): Geraniaceae: *California macrophylla, Erodium texanum, Geranium incanum, Geranium maderense, Geranium phaeum, Monsonia emarginata, Pelargonium cotyledonis*; Brassicaceae: *Capsella rubella*; Poaceae: *Oryza sativa*, *Sorghum bicolor*. Duplications of *ACC* were additionally identified in two Poaceae species—*Aegilops tauschii* and *Zea mays*—by performing BLAST searches against the genomes of these organisms on NCBI and Phytozome v13, respectively (Camacho et al., 2009; Goodstein et al., 2012).

All *ACC1* and *ACC2* sequences were included in a single file and aligned using the MAFFT *einsi* option (Katoh and Standley, 2013) in codon space using the *align_fasta_with_mafft_codon* subroutine in the sloan.pm Perl module (https://github.com/dbsloan/perl_modules). 5′ trimming was conducted according to the trimming performed in Rockenbach et al. (2016). Additional trimming of poorly aligned regions was performed manually in a codon-based manner.

### Compilation and curation of Clp core subunit amino acid and nucleotide sequences

To identify Clp core subunit amino acid sequences, a custom Python script (https://github.com/alissawilliams/Gene_duplication_ACCase_Clp/scripts/local_blast5.py) was used to reciprocally blast (blastp v2.2.29) *Arabidopsis thaliana* amino acid sequences against predicted protein sequences from each of 22 other angiosperm species in the dataset. These 22 species were the same set used in Williams et al. (2019) with the exclusion of *Silene latifolia* and *Silene noctiflora*, since Clp core subunit duplications have been previously studied in *Sileneae* (Rockenbach et al., 2016; Williams et al., 2021, 2019). This sampling was chosen to represent both the diversity of angiosperms and the range of rate variation in Clp complex evolution (Williams et al., 2019; see Table S3).

Compiled amino acid sequences for each subunit were aligned using the *einsi* option in MAFFT v7.222 (Katoh and Standley, 2013) and trimmed using GBLOCKS v0.91b (Castresana, 2000) with parameter *-b1* set to the default value of *-b2* and parameter -*b5* set to *h*. All alignments were examined manually to confirm homology. Sequences were also screened to prevent inclusion of multiple splice variants from a single gene. In cases where genomic data were used, only one transcript per gene was used. In cases where transcriptomic data were used, sequences were eliminated when alternative splicing was obvious (i.e. inclusion of an intron where the other sequence had a gap or variation only in one short piece of the transcript at either end). Catalytic site status and length were determined using the amino acid sequence data.

Nucleotide sequences for each identified Clp core subunit protein sequence were compiled from the corresponding CDS or transcript sequence file. For non-CDS sequences, ORFfinder (Wheeler et al., 2003) was used to identify the coding sequence. Compiled CDS sequences for each subunit were aligned with the MAFFT *einsi* option (Katoh and Standley, 2013) in codon space as above. 5′ and 3′ end trimming was performed manually in a codon-based manner.

### Generating constraint trees for the *ACC* and Clp subunit alignments for use in PAML

A constraint tree stipulates a fixed topology (branching order) that is used by a phylogenetic program (in this case, PAML) when calculating branch lengths. To generate a constraint tree for the *ACC* alignment, RAxML v8.2.12 (Stamatakis, 2014) was used on the trimmed nucleotide alignment with parameters *-m = GTRGAMMA, -p = 12345, -f = a, -x = 12345,* and *-# = 100*. The resultant topology confirmed that there were independent *ACC* duplications at the base of each family (Park et al., 2017; Rockenbach et al., 2016).

To construct constraint trees for Clp core subunits, each trimmed amino acid alignment was analyzed with ProtTest v3.4.2 (Darriba et al., 2011) to choose a model of sequence evolution. The top model based on the Bayesian Information Criterion was chosen for use in PhyML v3.3 (Guindon et al., 2010), which was run with 1000 bootstrap replicates and 100 random starts. The resultant phylogenetic trees were used to determine whether duplication events were lineage-specific or shared among species in the dataset. In almost all cases, paralogs from a single species were sister to one another in the trees, indicating lineage-specific duplications. There were a few cases in which paralogs from a single species were not sister to one another.

However, given low bootstrap support and the difficulty of resolving species relationships using a single gene with highly variable rates of evolution, we proceeded under the assumption that these duplications were lineage-specific as well. Thus, the constraint trees for each individual Clp core subunit were constructed using the known species tree (The Angiosperm Phylogeny Group et al., 2016), with duplications encoded as species-specific (mapped to terminal branches of the species tree).

### Running PAML for ACCase and Clp core subunit genes

For each alignment, PAML v4.9j (Yang, 2007) was used to infer *d*_N_/*d*_S_ values for all branches using the free ratios model (model = 1) and parameters *CodonFreq = 2* and *cleandata = 0*. Additionally, *model = 0* and *model = 2* runs were conducted for all alignments, again using *CodonFreq = 2* and *cleandata = 0*. The *model = 0* runs forced all branches to have the same *d*_N_/*d*_S_ ratio, while the *model = 2* runs allowed different *d*_N_/*d*_S_ values for specified groups of branches.

For the *ACC* alignment, one *model = 2* run was conducted with plastid-targeted branches as the foreground. The resultant tree had one *d*_N_/*d*_S_ value for plastid-targeted (*ACC2*) branches and a second *d*_N_/*d*_S_ value for cytosol-targeted (*ACC1*) branches (including all internal pre-duplication branches). This output was compared with the *model = 0* run to determine whether allowing two *d*_N_/*d*_S_ ratios (one for each of those groups) was a better fit to the data than allowing just a single *d*_N_/*d*_S_ value. For the Clp subunit alignments, *model = 2* was used twice. In the first run, all terminal branches (and in the case of two subunits, internal post-duplication branches) were designated as the foreground. In the second run, there were three classes of branches, where all branches were categorized the same as in the first run except that post-duplication branches (internal or terminal) were placed in a third category. The three-partition and two-partition models were compared to determine whether allowing an additional *d*_N_/*d*_S_ ratio for post-duplication branches was a better fit to the data than just separating terminal from internal branches. The models were compared using likelihood ratio tests.

For the *ACC* alignment, a branch-site test (Yang, 2007; Yang and Nielsen, 2002) was also conducted to test for evidence of positive selection on branches for plastid-targeted genes, which were set as the foreground branches for this analysis. A null model and an alternative model both used the parameters *model = 2, Nssites = 2, CodonFreq = 2*, and *cleandata = 0*. The alternative model otherwise used all default values, while the null model additionally used *fix_omega = 1* and *omega = 1*. The models were compared using a likelihood ratio test.

### Running HyPhy for *ACC*

In addition to running a PAML branch-site test on the *ACC* alignment (Yang, 2007; Yang and Nielsen, 2002), tests for positive and relaxed selection were implemented in HyPhy v2.5.32 (Kosakovsky Pond et al., 2020). Positive selection was tested for using the aBSREL and BUSTED methods (Murrell et al., 2015; Smith et al., 2015). The RELAX method was used to test for relaxed vs. intensified selection (Wertheim et al., 2015). As with the PAML runs, the constraint tree used for HyPhy methods had the branches separated into two categories (*ACC1* and *ACC2*).

### Comparisons between *ACC1* and *ACC2* genes

To compare *d*_N_ and *d*_S_ between genes encoding cytosolic-targeted proteins and genes encoding plastid-targeted proteins (*ACC1* and *ACC2*, respectively), a mean root-to-tip distance was calculated for each family in the tree. The base of each duplication event was used as the root for each family. For both *d*_N_ and *d*_S_, the four mean distances for *ACC1* were compared to those of *ACC2* using a paired t-test in R. Because of the *a priori* prediction that retargeting to the plastid would be associated with accelerated protein sequence evolution, a one-sided test (*ACC2* > *ACC1*) was used for *d*_N_, while a two-sided test was used for *d*_S_.

### Fisher’s exact test on Clp subunit paralogs

Using the output from the free ratios (model = 1) PAML runs, a two-sided Fisher’s exact test was used to test for asymmetry in the ratio of the estimated numbers of nonsynonymous substitutions (N) and synonymous substitutions (S) (Pegueroles et al., 2013). Nonsynonymous and synonymous substitution estimates were entered into the *fisher.test()* function in R with default parameters. For each pair of duplicates, a test between paralog 1 and paralog 2 was performed (**Figure 1**). If the paralogs were found to be evolving symmetrically, their combined numbers of substitutions were compared to those of the ancestral branch (**Figure 1**). If the paralogs were found to be evolving asymmetrically, each one was compared individually against the ancestral branch (**Figure 1**). The four cases in which there were more than two species-specific paralogs (*Soja max* and *Gossypium raimondii CLPP5*; *Musa acuminata* and *Vitis vinifera CLPR4*) were excluded from this analysis.

**Figure 1:**
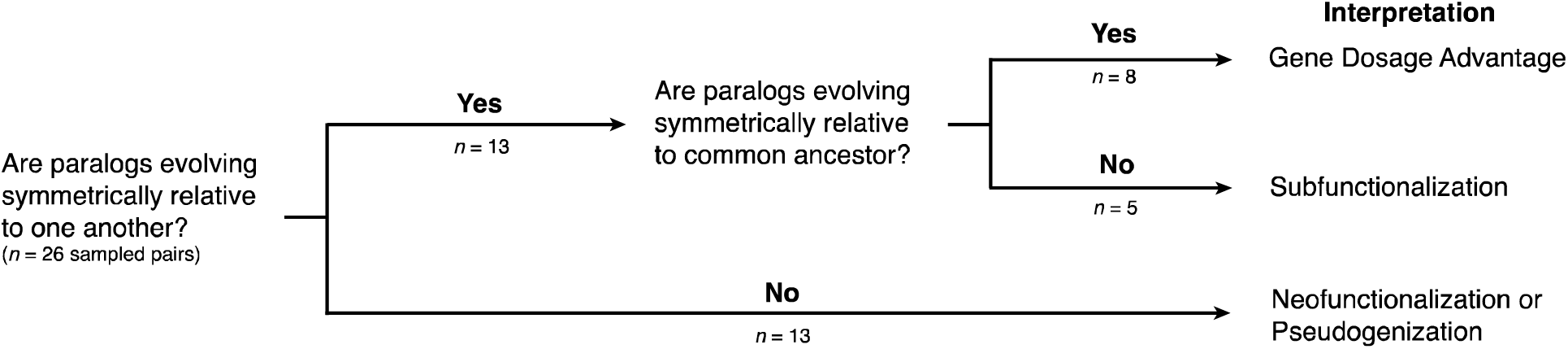
Expectations under different post-duplication models. For each pair of paralogs (n = 26), we first determined whether they were evolving symmetrically relative to one another using N and S estimates from PAML output. Paralogs evolving asymmetrically are predicted to represent neofunctionalization or pseudogenization events. For paralogs evolving symmetrically (n = 8), combined N and S values were compared to those of the immediate ancestor branch. Pairs evolving symmetrically relative to the common ancestor (n = 8) are predicted to represent gene dosage advantage while those evolving asymmetrically relative to the common ancestor (n = 5) are predicted to represent subfunctionalization.

### Data availability

Scripts, untrimmed and trimmed alignments, PAML output, and HyPhy output are provided for both *ACC* and Clp subunits at https://github.com/alissawilliams/Gene_duplication_ACCase_Clp.

## Results

### Plastid-targeted ACCases evolve more rapidly than cytosol-targeted ACCases across angiosperms

Across the sampled clades (Geraniaceae, Caryophyllaceae, Brassicaceae, and Poaceae), nearly all *ACC2* genes (which encode plastid-targeted proteins) have higher *d*_N_/*d*_S_ values than their *ACC1* counterparts (which encode cytosol-targeted proteins) (**Figure 2, Figure S1**). The single-partition model assigned all branches a *d*_N_/*d*_S_ value of 0.1266, while the two-partition model assigned *ACC1* branches a value of 0.0883 and *ACC2* branches a value of 0.1936 (χ^2^ = 466.84, p << 0.0001). This pattern is true for both terminal and internal branches. The increase in in *d*_N_/*d*_S_ ratios in *ACC2* branches is generally driven by increases in *d*_N_ rather than reductions in *d*_S_ (t = 4.48, p = 0.01 for *d*_N_; t = 0.72, p = 0.5249 for *d*_S_; **Figure 2, Figure S2**), suggesting changes in selective pressure.

**Figure 2:**
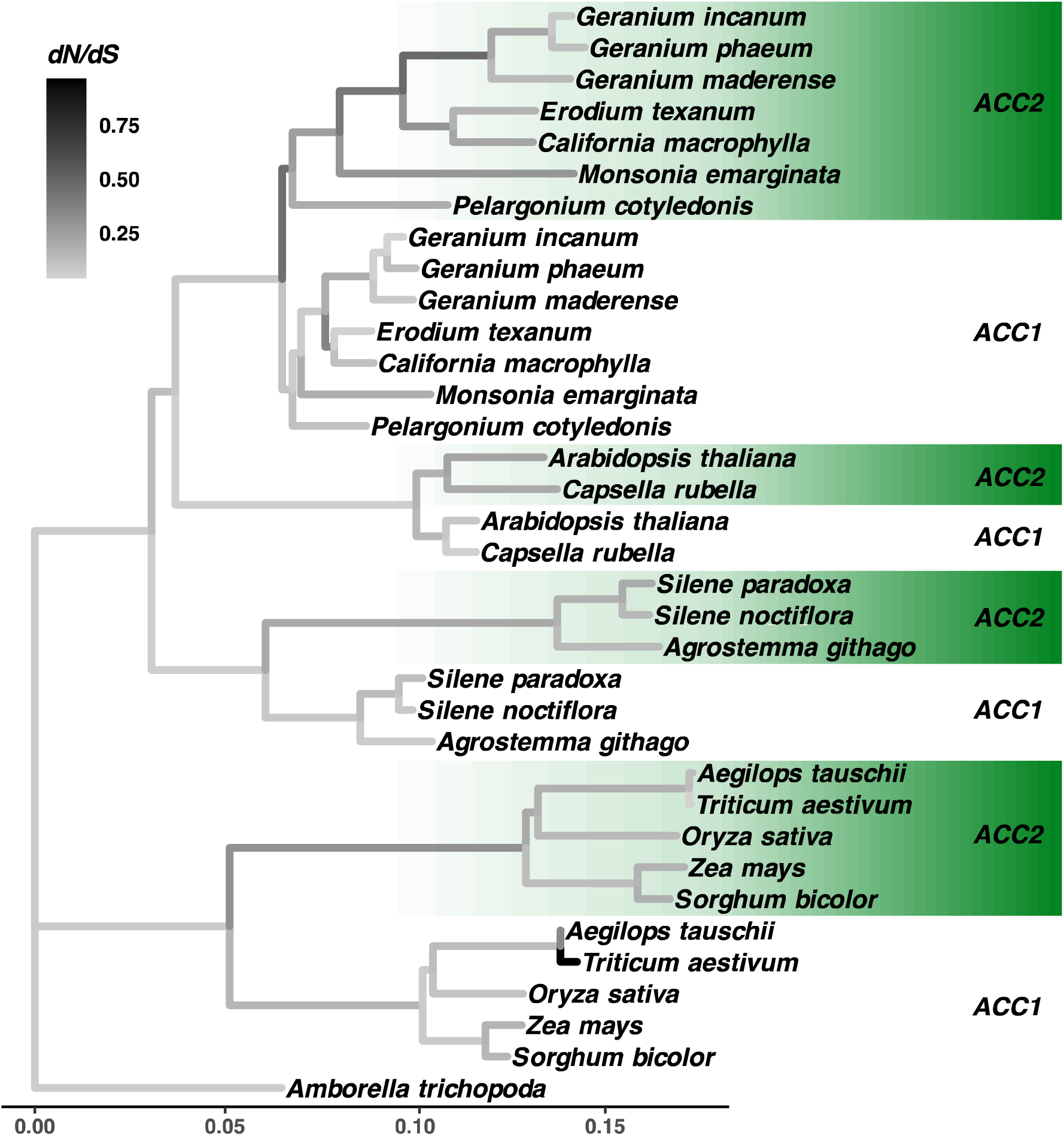
*ACC* genes across the Brassicaceae, Caryophyllaceae, Geraniaceae, and Poaceae, with the single copy of *ACC* in *Amborella trichopoda* as an outgroup. Branch lengths represent *d_N_* values and branch colors represent *d*_N_/*d*_S_ ratios.

Using a branch-sites test in PAML (Yang, 2007), we did not find a significant signature of positive selection spanning the alignment (χ^2^ = 0, p = 1), although there were multiple individual sites found to be under positive selection (**Table S1**). Two HyPhy methods found limited, though significant, evidence for positive selection—the aBSREL run (Smith et al., 2015) detected one branch under positive selection (p = 0.04) and the BUSTED run (Murrell et al., 2015) assigned 0.12% of sites in foreground (*ACC2*) branches to the positive selection class relative to 0.05% of sites in background (*ACC1*) branches (p = 0.0026). The HyPhy RELAX method (Wertheim et al., 2015) found significant evidence for relaxed selection in the *ACC2* branches relative to the rest of the tree (K = 0.09, p <<0.001).

**Table 1:** Differences in evolutionary rates between duplicated and non-duplicated plastid Clp core subunits. Reported *p-*values are based on likelihood ratio tests for 2-partition vs. 3-partition PAML models (see Materials and Methods). Log-likelihood (lnL) values are reported for each model.

| Subunit | lnL 2-partition model | lnL 3-partition model | *p*-value | Class with higher $d_N/d_S$ |
| --- | --- | --- | --- | --- |
| *CLPP3* | -10058.01 | -10047.93 | 7.12e-06 | Duplicated |
| *CLPP4* | -9606.41 | -9552.26 | <1.00e-10 | Duplicated |
| *CLPP5* | -8121.80 | -8070.90 | <1.00e-10 | Duplicated |
| *CLPP6* | -7697.62 | -7691.31 | 3.82e-04 | Non-duplicated |
| *CLPR1* | -14534.75 | -14517.85 | 6.07e-09 | Duplicated |
| *CLPR2* | -12018.69 | -12002.74 | 1.63e-08 | Duplicated |
| *CLPR3* | -10556.37 | -10556.05 | 0.42 | n/a |
| *CLPR4* | -10395.60 | -10337.35 | <1.00e-10 | Duplicated |

### Characterizing ongoing duplication of nuclear-encoded Clp core subunit genes in angiosperms

Of the 23 angiosperm species in our dataset, 11 had one or more duplications of nuclear genes encoding Clp core subunits, and all eight of these genes were duplicated in at least one species (**Figure 3**). Most of these duplications were represented by two paralogs, but in four cases, we identified more than two paralogs for a particular subunit in a particular species. For *CLPP5*, *Soja max* and *Gossypium raimondii* have five and seven copies, respectively, and for *CLPR4*, both *Musa acuminata* and *Vitis vinifera* have four copies.

**Figure 3:**
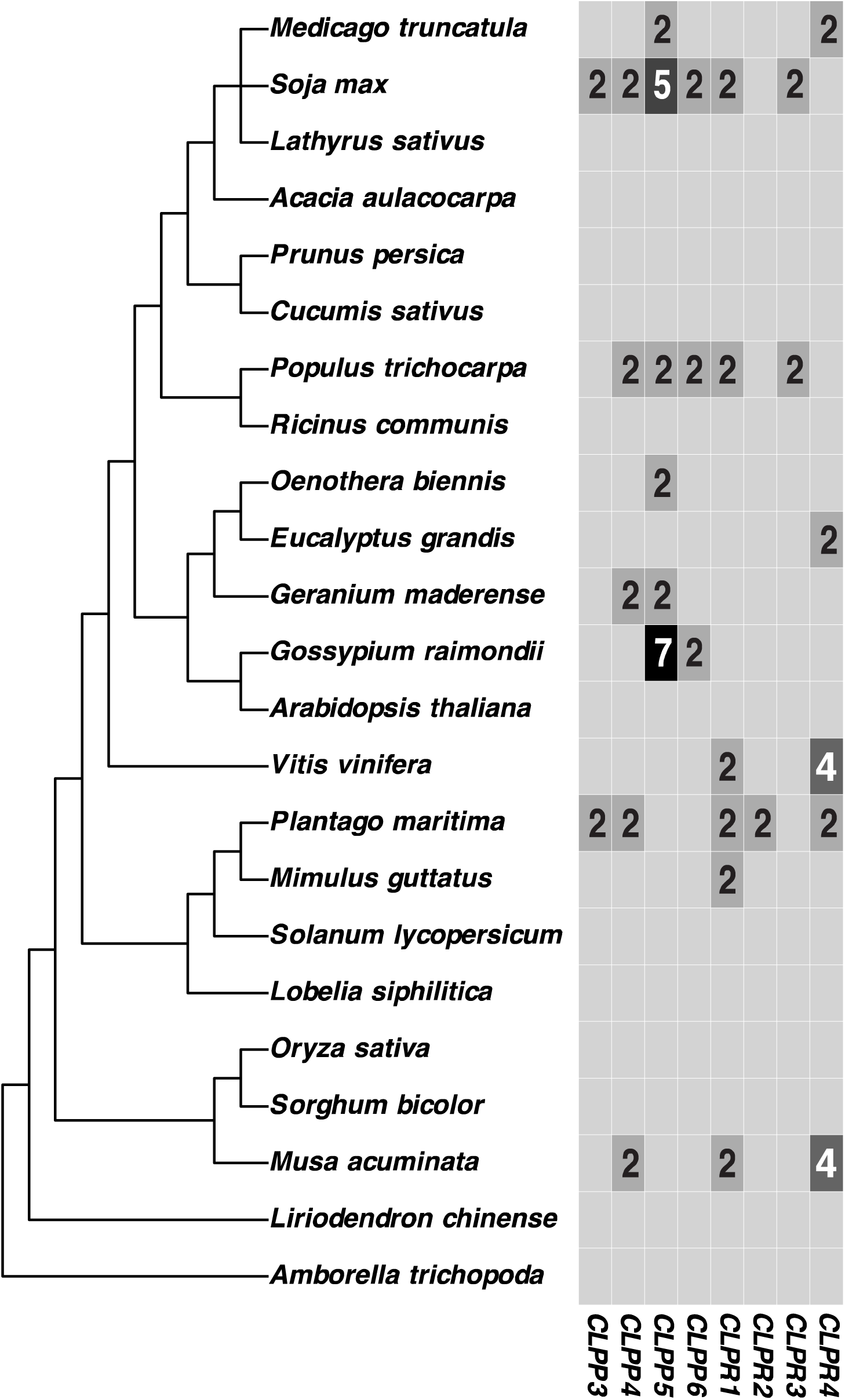
Copy numbers of the nuclear-encoded subunits of the plastid Clp core across angiosperms. Boxes without numbers indicate single-copy genes.

*Soja max* had duplications of the largest number of subunits (six of eight), followed by *Plantago maritima* and *Populus trichocarpa* with duplications of five subunits. Of the 11 species with duplications, *Eucalyptus grandis* and *Oenothera biennis* were the only species that had duplications of just one subunit. Across subunits, *CLPP5* had the highest number of paralogs (37 in 23 species) and *CLPR2* had the lowest (24 in 23 species).

In total, we identified 72 gene copies of Clp core subunits resulting from duplication events, including 40 catalytic subunits (*CLPP3*-*CLPP6*) (**Figure 3**). Of the 40 paralogs of catalytic subunits, we found evidence of loss of one or more catalytic sites in multiple genes (**Table S2**). Across all 72 paralogs, we also found evidence of truncation of multiple different gene copies (including some with catalytic site loss) (**Table S2, Table S3**).

**Table 2:** *p*-values for asymmetries between paralogs and between paralogs and their common ancestor.

| Species | Gene | Paralog 1 vs.<br>paralog 2 | Paralog 1 vs.<br>ancestor | Paralog 2 vs.<br>ancestor | Paralogs 1+2<br>vs. ancestor |
| --- | --- | --- | --- | --- | --- |
| *P. maritima* | *CLPP3* | 0.83 |  |  | 1.25e-04* |
| *S. max* | *CLPP3* | 1 |  |  | 0.20 |
| *P. maritima* | *CLPP4* | 1.28e-07 | 1.07e-09* | 0.71 |  |
| *M. acuminata* | *CLPP4* | 0.04 | 0.15 | 2.12e-04* |  |
| *S. max* | *CLPP4* | 0.57 |  |  | 1 |
| *P. trichocarpa* | *CLPP4* | 3.31e-04 | 3.44e-12* | 1.76e-13* |  |
| *G. maderense* | *CLPP4* | 0.08 |  |  | 1.72e-03* |
| *M. truncatula* | *CLPP5* | 1 |  |  | 3.61e-05* |
| *P. trichocarpa* | *CLPP5* | 1 |  |  | 0.04^ |
| *O. biennis* | *CLPP5* | 9.12e-3 | 0.10 | 5.37e-4* |  |
| *G. maderense* | *CLPP5* | 1.71e-04 | 0.42 | 6.23e-4* |  |
| *S. max* | *CLPP6* | 0.27 |  |  | 1 |
| *P. trichocarpa* | *CLPP6* | 0.46 |  |  | 1 |
| *G. raimondii* | *CLPP6* | 0.71 |  |  | 1 |
| *M. acuminata* | *CLPR1* | 2.20e-16 | 0.82 | 2.20e-16* |  |
| *S. max* | *CLPR1* | 0.47 |  |  | 0.05^ |
| *P. trichocarpa* | *CLPR1* | 0.77 |  |  | 0.42 |
| *V. vinifera* | *CLPR1* | 4.45e-3 | 0.17 | 1.38e-09* |  |
| *M. guttatus* | *CLPR1* | 0.05 | 0.59 | 4.70e-04* |  |
| *P. maritima* | *CLPR1* | 4.70e-05 | 0.53 | 2.54e-05* |  |
| *P. maritima* | *CLPR2* | 0.01 | 0.18 | 1.63e-06* |  |
| *S. max* | *CLPR3* | 1 |  |  | 0.55 |
| *P. trichocarpa* | *CLPR3* | 0.34 |  |  | 0.35 |
| *M. truncatula* | *CLPR4* | 2.29e-03 | 0.21 | 0.11 |  |
| *E. grandis* | *CLPR4* | 2.51e-05 | 0.68 | 5.42e-11* |  |
| *P. maritima* | *CLPR4* | 9.73e-05 | 9.53e-04 | 4.17e-14* |  |

\* denotes that paralog(s) has/have significantly higher evolutionary rate than ancestor branch
^ denotes that paralog(s) has/have significantly lower evolutionary rate than ancestor branch

### Recent paralogs of Clp core subunits tend to have higher rates of protein sequence evolution than their single-copy counterparts

Out of the eight nuclear-encoded Clp core subunit trees (Figures S3**-**S10), seven showed statistically significant differences between a model that allowed for different *d*_N_/*d*_S_ rates in gene duplicates vs. single-copy genes (the three-partition model) and one that forces the same *d*_N_/*d*_S_ rate on these two types of branches (the two-partition model) based on an uncorrected significance threshold of *p* = 0.05. (**Figure 4, Table 1**). In six of those cases, duplicated terminal branches had a higher *d*_N_/*d*_S_ rate than non-duplicated terminal branches, while in the remaining case, the reverse was true. We separated internal branches from terminal branches to account for the fact that terminal branches will, on average, have higher *d*_N_/*d*_S_ estimates than internal branches because selection has had more time to act on older deleterious mutations (Hasegawa et al., 1998; Ho et al., 2005). Further, terminal branches represent both interspecific divergence and intraspecific polymorphism, which is important because the latter inflates evolutionary rate calculations (Ho et al., 2005; Moilanen and Majamaa, 2003; Nielsen and Weinreich, 1999).

**Figure 4:**
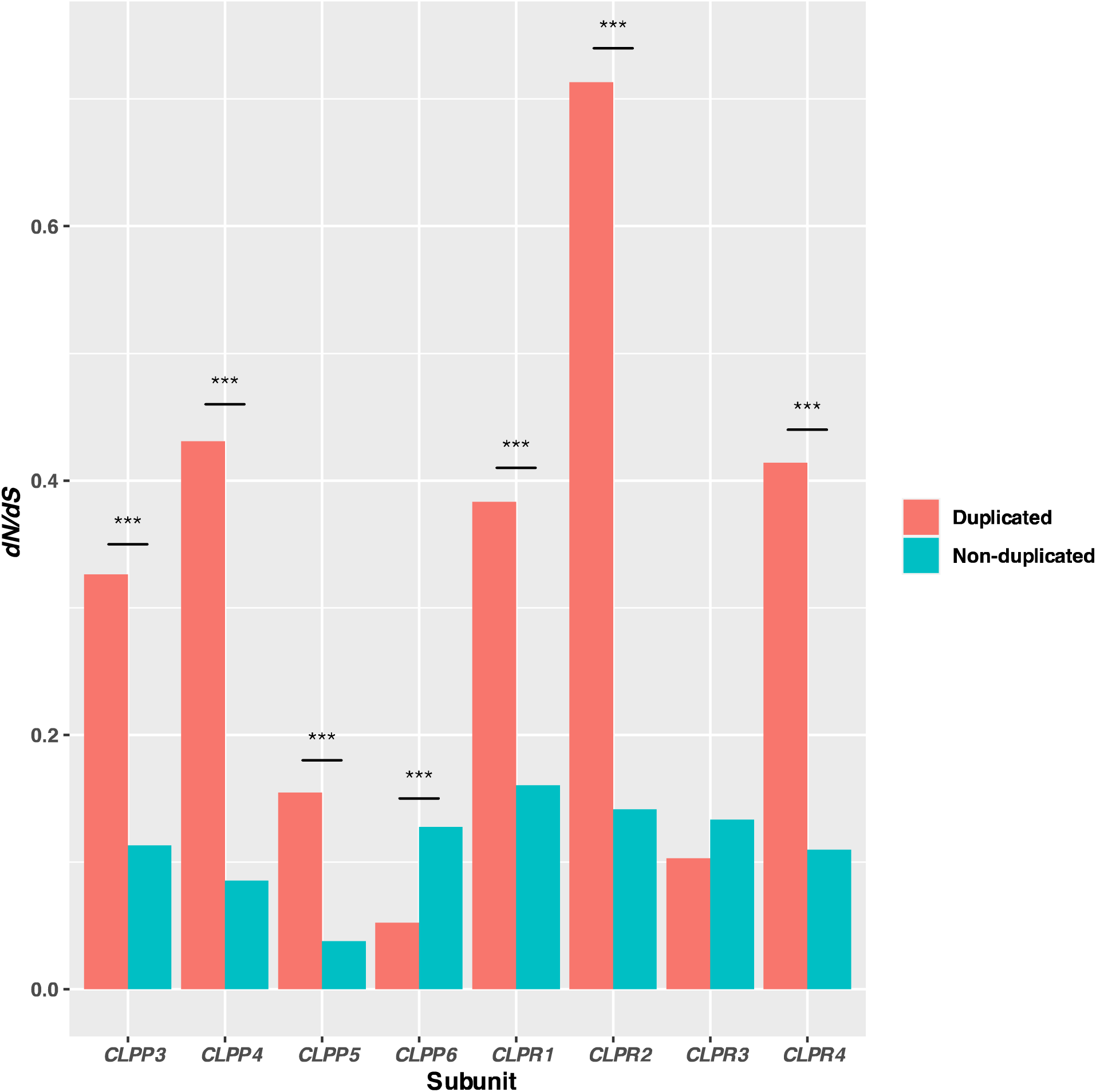
*d*_N_/*d*_S_ ratios of duplicated and non-duplicated plastid Clp core subunits across angiosperms. The values were calculated using a PAML branch test with three groups, where each group was assigned its own *d*_N_/*d*_S_ value: non-duplicated terminal branches, duplicated terminal branches (and in the cases of *CLPP5* and *CLPR4*, internal post-duplication branches), and internal branches. Significant differences (p<<0.001) are indicated with ***.

We also compared the ratios of estimated nonsynonymous substitutions (N) to estimated synonymous substitutions (S) between paralogs as well as between paralogs and their common ancestor, again using an uncorrected significance threshold of p = 0.05 (**Table 2**). Of the 26 pairs of paralogs, 13 (50%) showed statistically significant differences in N/S ratios relative to each another. In 10 (77%) of those cases, only one paralog had a significantly different N/S ratio than that of the common ancestor (and in all 10 of those cases, that paralog had a higher N/S ratio than that of the common ancestor). Of the 13 pairs with N/S ratios that were not statistically different from one another, five (38%) had different N/S ratios relative to the common ancestor. In three of those cases, the combined paralog N/S ratio was significantly higher than that of the ancestor.

## Discussion

### Accelerated evolution of duplicated *ACC* genes in multiple clades of flowering plants

We found that branches of our *ACC* tree representing proteins targeted to the plastid had statistically significantly higher *d*_N_/*d*_S_ values than branches representing proteins targeted to the cytosol (**Figure 2, Figure S1**), consistent with the predictions under neofunctionalization. These results were based on a trimmed alignment lacking the target peptide, which we excluded because target peptides exhibit fast rates of evolution and reduced constraints on primary amino acid sequence (Bruce, 2001, 2000; Jarvis, 2008). Thus, our results show that *ACC* genes encoding proteins retargeted to the plastid are undergoing evolutionary rate increases unrelated to the target peptide, suggesting that other functional domains are also evolving rapidly.

Retargeting of the cytosolic, homomeric ACCase protein to the plastid is somewhat unexpected given that a heteromeric ACCase complex already exists in plastids. Whether the retargeted homomeric ACCases functionally replaces or coexists with the heteromeric version appears to vary across clades. In some angiosperm groups, the two complexes coexist, including in *Arabidopsis thaliana* and likely in other members of the Brassicaceae (Babiychuk et al., 2011; Rousseau-Gueutin et al., 2013). In other clades, the homomeric ACCase has replaced the heteromeric version, as was reported in the Poaceae (Konishi and Sasaki, 1994). The duplication found in *Silene noctiflora* and *Silene paradoxa* may also represent a replacement event given that both species lack at least one heteromeric ACCase gene each, where *S. noctiflora* lacks all of them (Rockenbach et al., 2016). In some cases, including *Monsonia emarginata* in the Geraniaceae, the plastid-encoded *accD* gene of the heteromeric complex has been transferred to the nuclear genome, again suggesting that the heteromeric version is still functional (Park et al., 2017; Rousseau-Gueutin et al., 2013). In the gymnosperm species *Gnetum ula*, there are two nuclear-encoded copies of *accD* whose protein products are differentially targeted, suggesting that in some species of plants, there are multiple types of heteromeric ACCase (Sudianto and Chaw, 2019). The contrasting histories of replacement vs. coexistence may mean that duplicates in different clades are evolving under different selection regimes.

Variation in post-duplication fates could confound tests of selection conducted across the entire *ACC* tree. Using PAML and HyPhy (Murrell et al., 2015; Smith et al., 2015; Wertheim et al., 2015; Yang, 2007), we tested for positive selection and relaxed selection in *ACC2* genes relative to *ACC1* genes, both of which can contribute to increased rates of protein sequence evolution. The results were mixed; there is some evidence for relaxed selection across all *ACC2* branches as well as for positive selection in a small number of branches and sites (**Table S1**). Across the four families in our sample, the smallest ratio between mean *ACC2 d*_N_ and mean *ACC1 d*_N_ was found in the Poaceae (1.5 vs. 2.2-2.6 for the other three families). Since the heteromeric ACCase is completely absent in the Poaceae (Konishi and Sasaki, 1994), we would expect stronger purifying selection on the plastid homomeric ACCase in this clade compared to clades in which the two versions coexist. Thus, these results are consistent with the hypothesis that relaxed selection is contributing to rate accelerations and that there is greater relaxation of selection when homomeric and heteromeric ACCases functions redundantly in the plastid, though the evidence is still limited. The potential for positive selection on retargeted ACCases is intriguing given that these proteins are thought to perform the same function as the ancestral protein; it is possible that retargeted proteins are adapting to specific biochemical and/or osmotic conditions within the new destination. Increased evolutionary rates after subcellular retargeting have been previously noted, though we do not fully understand their underlying causes (Byun-McKay and Geeta, 2007; Marques et al., 2008).

### Ongoing duplication of nuclear-encoded Clp core subunit genes is common in angiosperms

Across green plants, duplication of the plastid-encoded Clp core subunit gene *clpP1* has only been found in a handful of lineages (Williams et al., 2019). While other studies have identified recent duplications of nuclear-encoded Clp core subunit genes (Rockenbach et al., 2016; Williams et al., 2021), our work shows that duplications of these nuclear-encoded subunits are pervasive across angiosperms (**Figure 3**). Because we used a mix of transcriptomic and genomic data, we took into consideration the possibility of misidentifying transcript variants as paralogs but our use of primary transcripts only and manual curation to remove hits that appeared to be splice variants (see Materials and Methods) minimizes the risk of this type of error.

The prevalence of whole genome duplication in plants may partially explain the prevalence of Clp core subunit duplication (Clark and Donoghue, 2018; De Bodt et al., 2005; del Pozo and Ramirez-Parra, 2015; Flagel and Wendel, 2009; Panchy et al., 2016; Wendel et al., 2018). For instance, *Soja max* is a partially diploidized tetraploid, meaning that this lineage underwent a polyploidization event very recently and has only just started the subsequent process of genome reduction (Shultz et al., 2006). *Soja max* had the largest number of duplicated subunits across our sample, which is consistent with this history of whole genome duplication. Similarly, *Populus trichopoda*, which tied for the second largest number of duplicated subunits, only recently underwent genome reduction after whole genome duplication (Tuskan et al., 2006). In these cases, we may simply be observing the short-term effects of polyploidization prior to returning to a single copy of each of these genes.

### Possible subfunctionalization in recent paralogs of Clp core subunits

Clp core subunit ratios have been studied in *Arabidopsis thaliana* (Olinares et al., 2011a). The core consists of two rings—a ClpP1/ClpR1-4 ring with a 3:1:1:1:1 subunit ratio, respectively, and a ClpP3-6 ring with a 1:2:3:1 subunit ratio, respectively (Olinares et al., 2011a). Despite the high degree of structural similarity amongst the plastid Clp core subunits, core composition (i.e. the number of each type of core subunit) does not appear to vary in *A. thaliana* (Olinares et al., 2011a; Peltier et al., 2004). Due to the stability of subunit interactions in *A. thaliana*, Clp complexes in other angiosperms are typically assumed to have the same ratios of core subunits, but our results suggest that varied numbers of core subunit paralogs may lead to varied stoichiometry across species. Additional work has shown that loss of catalytic activity in ClpP5 (present in three copies in *A. thaliana*) is lethal while loss of catalytic activity in ClpP3 (present in one copy in *A. thaliana*) is tolerated, suggesting that core subunit composition may be flexible given a threshold number of catalytic subunits (Liao et al., 2018).

Given that subfunctionalization has likely played a major role in plastid Clp complex evolution, we were particularly interested in whether we could identify subfunctionalization after more recent duplication events. Taken to an extreme, subfunctionalization would involve having one gene for each of the 14 core subunits, which would lead to further expansion of the typical nine core subunit genes. The total number of core subunits after including recent paralogs and the plastid-encoded ClpP1 was less than 14 in most species. *Musa acuminata*, *Plantago maritima*, and *Populus trichocarpa* had 14 each, *Soja max* had 17, and *Gossypium raimondii* had 15 (**Figure 3)**. The numbers larger than 14 were driven in both cases by multiple paralogs of *CLPP5*, with five and seven copies, respectively. ClpP5 has the largest number of subunits stoichiometrically among the eight nuclear-encoded subunits, so the fact that the two largest numbers of paralogs were both found in *CLPP5* could potentially suggest that some species are moving toward a 1:1 relationship between genes and core subunits. However, this explanation is not supported by other evidence. For example, the other cases of >2 paralogs were found for *CLPR4*, which encodes a protein that is present in just a single copy in the core in *Arabidopsis*. Further, it is not clear that all of these paralogs are capable of producing functional proteins given truncations and loss of catalytic sites (**Table S2, Table S3**).

We tested for signatures of subfunctionalization by looking at asymmetry in N/S ratios. Under subfunctionalization, we would expect paralogs to evolve at symmetric rates relative to one another but asymmetrically relative to their common ancestor (Pegueroles et al., 2013). We found five of these cases in our dataset: *Plantago maritima CLPP3*, *Geranium maderense CLPP4, Medicago truncatula CLPP5, Populus trichocarpa CLPP5,* and *Soja max CLPR1* (**Table 2**). In cases of subfunctionalization, we would expect the paralogs to evolve more quickly than the common ancestor because of relaxed selection due to their more limited functional roles, which was only the case for the former three. In those three cases, the evidence is consistent with subfunctionalization, particularly given that all six involved paralogs are full length. Further, the *P. maritima CLPP3* paralogs share the same substitutions in all three catalytic sites, which indicates duplication after the loss of catalytic activity, and the *G. maderense CLPP4* and *M. truncatula CLPP5* paralogs all have fully retained catalytic triads (**Table S2**).

### Possible pseudogenization or neofunctionalization in recent paralogs of Clp core subunits

Predictions about evolutionary rates under neofunctionalization are similar to predictions under the degeneration/gene loss model—one paralog will maintain the ancestral evolutionary rate while the other undergoes evolutionary rate acceleration (Hahn, 2009; Pegueroles et al., 2013; Zhang, 2003). Previous work in this complex has shown that even ClpP1 subunits demonstrating massive accelerations in evolutionary rate can still be functional, meaning that high evolutionary rates alone do not necessarily indicate pseudogenization (Barnard-Kubow et al., 2014; Williams et al., 2019, 2015). Other sequence features can help us differentiate between pseudogenization and neofunctionalization. For example, truncation of a sequence can be evidence that it is no longer producing a functional protein; additionally, for ClpP subunits, loss of catalytic sites may also be an indication of degeneration/pseudogenization (though there may be exceptions, including the *P. maritima CLPP3* paralogs mentioned above). In our dataset, the paralogs of *Musa acuminata CLPP4* and *P. trichocarpa CLPP4* follow these patterns (**Table 2**). In each of these pairs, the paralogs are evolving asymmetrically, and the paralog with a faster rate of evolution is truncated and lacking all three catalytic sites, suggesting loss of function (**Table S2**). Another example of probable pseudogenization is found for the second copy of *M. acuminata CLPR1*. This paralog was annotated as two separate genes due to an internal stop codon, which would lead to a truncation in the resultant protein.

As for neofunctionalization, there are other cases in our dataset where paralogs with asymmetric N/S ratios both have retained catalytic sites and are full length (for instance, *Geranium maderense CLPP5* and *Oenothera biennis CLPP5*). There are no known instances of retargeting of plastid Clp core subunits; thus, evolutionary drivers of neofunctionalization of duplicated subunits are unknown. It is possible that neofunctionalization in this complex could involve recruiting additional interacting proteins—the ClpT proteins, for instance, are involved in assembly of the core and are a recently evolutionary innovation specific to green plants (Colombo et al., 2014; Kim et al., 2015; Nishimura and van Wijk, 2015; Sjögren and Clarke, 2011). Additionally, ongoing work has identified potential new adapter proteins in the plastid Clp complex (Montandon et al., 2019; Nishimura et al., 2015). Another possibility is tissue-specific expression of paralogs, which has not been documented in the Clp complex but has been identified in mitochondrial complexes (Boss et al., 1997; Guerrero-Castillo et al., 2017; Sinkler et al., 2017).

### Possible retention of Clp core paralogs under the gene dosage advantage hypothesis

We additionally identified eight cases of paralogs with symmetric N/S ratios where those ratios are also symmetric relative to the common ancestor (**Table 2**). Under our initial predictions, these would represent paralogs retained under the gene dosage advantage hypothesis (Hahn, 2009; Ohno, 1970; Pegueroles et al., 2013; Zhang, 2003). Of these eight paralog pairs, four are from *Soja max* (which had six total pairs of paralogs), and three are from *Populus trichocarpa* (which had five total pairs of paralogs). As described above, both of these species are in the process of rediploidization after a recent whole genome duplication (Shultz et al., 2006; Tuskan et al., 2006). It is possible that these results reflect the fact that the gene duplications happened so recently that the paralogs have not had time to diverge. This possibility is further supported given that the estimates of numbers of substitutions for many of these paralogs were so low that there was virtually no power to detect significant asymmetry.

### Alternative hypotheses and future directions

While we based our analyses on established expectations for evolutionary rates under different post-duplication fates (gene dosage advantage, neofunctionalization, and subfunctionalization), other work has challenged the universality of these predictions. He and Zhang (2005) outline the subneofunctionalization model, in which gene duplicates undergo rapid subfunctionalization followed by prolonged neofunctionalization. Asymmetric evolutionary rates are often assumed to be the result of either neofunctionalization or degeneration, but subfunctionalization can also occur in an asymmetric fashion (He and Zhang, 2005). This hypothesis could relate to some of our results; cases of asymmetric evolutionary rates could be due to subfunctionalization rather than neofunctionalization. Additionally, functional constraint can also exist under neofunctionalization, leading to lower substitution rates and possibly symmetric rates of evolution, meaning that symmetrically evolving paralogs could represent cases of neofunctionalization rather than subfunctionalization or gene dosage advantage (He and Zhang, 2005).

Regardless, our results demonstrate that post-duplication evolutionary fates of paralogs vary widely across clades, even when the same genes are involved. Duplications of the homomeric ACCase complex gene (*ACC*) and subsequent retargeting of one protein to the plastid have been previously reported (Babiychuk et al., 2011; Konishi and Sasaki, 1994; Park et al., 2017; Parker et al., 2014; Rockenbach et al., 2016; Schulte et al., 1997). Our results show that the retargeted duplicates almost universally have increased *d*_N_/*d*_S_ rates (**Figure 2, Figure S1**). As for plastid Clp core subunit duplications, duplication has clearly shaped this complex over the course of Viridiplantae evolution. We provide evidence of all possible post-duplication routes of recent paralogs amongst the different subunits and different species in our dataset. Overall, our results provide compelling evidence that subunit ratios and stoichiometry may be dynamic across angiosperm lineages. Isolation of plastid Clp complexes and analyses of subunit composition have been performed in a handful of species (Moreno et al., 2017; Olinares et al., 2011a; Williams et al., 2019); future work could determine these compositions in other angiosperms, including those that have undergone recent gene duplications. Our work demonstrates that gene duplication has been and continues to be an important force in plastid evolution.

## Acknowledgements

We thank two anonymous reviewers for their helpful comments on an earlier version of this manuscript. We also thank Patricia Bedinger, Mark Stenglein, Rachel Mueller, and Marinus Pilon for their feedback on this project and manuscript. This work was supported by a National Science Foundation (NSF) grant (MCB-1733227), graduate fellowships from NSF (DGE-1321845) and the National Institutes of Health (T32-GM132057), and the Wolves to Rams undergraduate research program (NSF Grant Numbers 1930150 and 19300092, NIH Grant Number 1T34GM137861-01).

---

**Figure S1.**
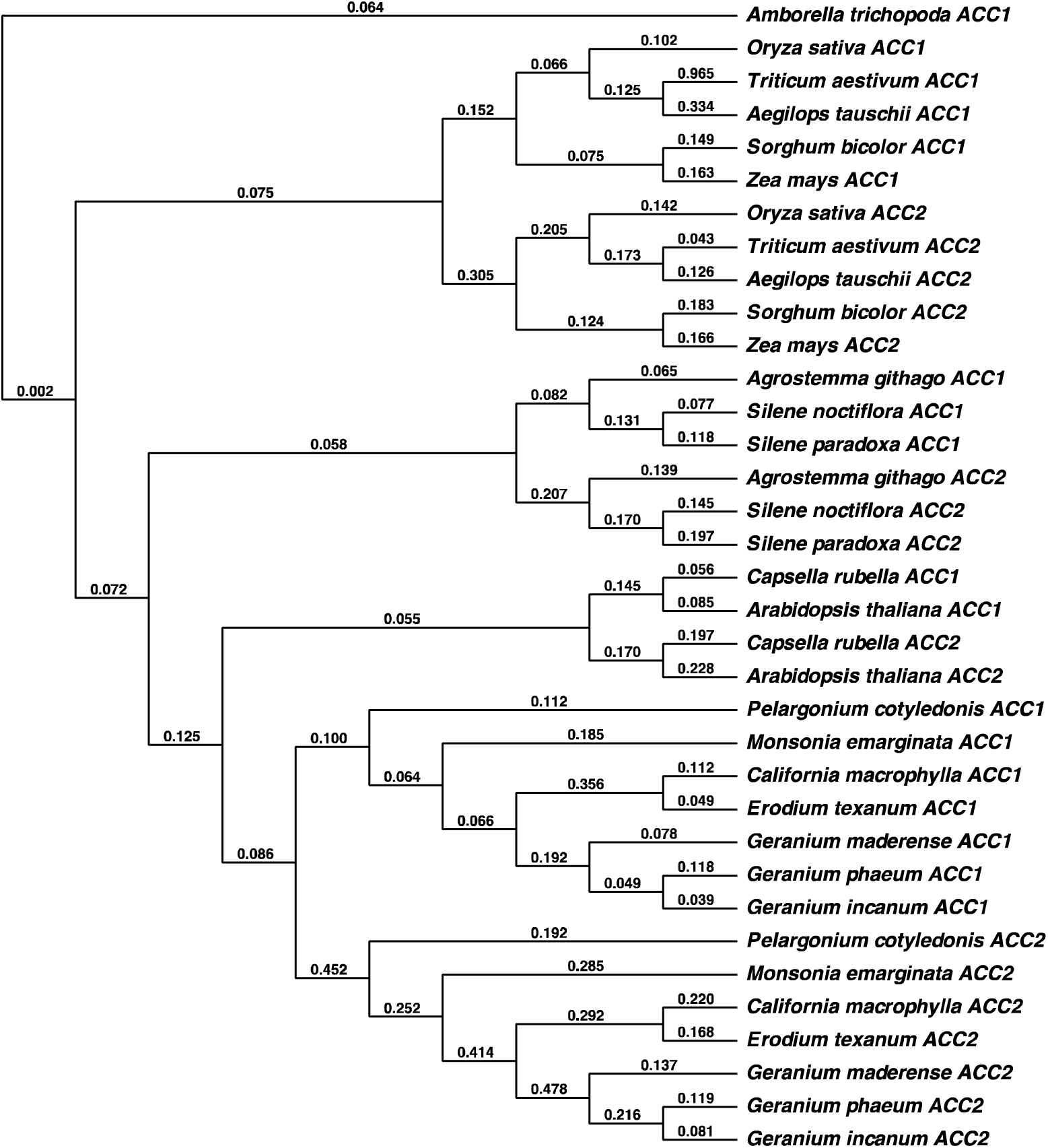
*ACC* tree. Branch labels are *d_N_*/*d_S_* values. *ACC1* represents cytosolic-targeted genes while *ACC2* represents plastid-targeted genes.

**Figure S2.**
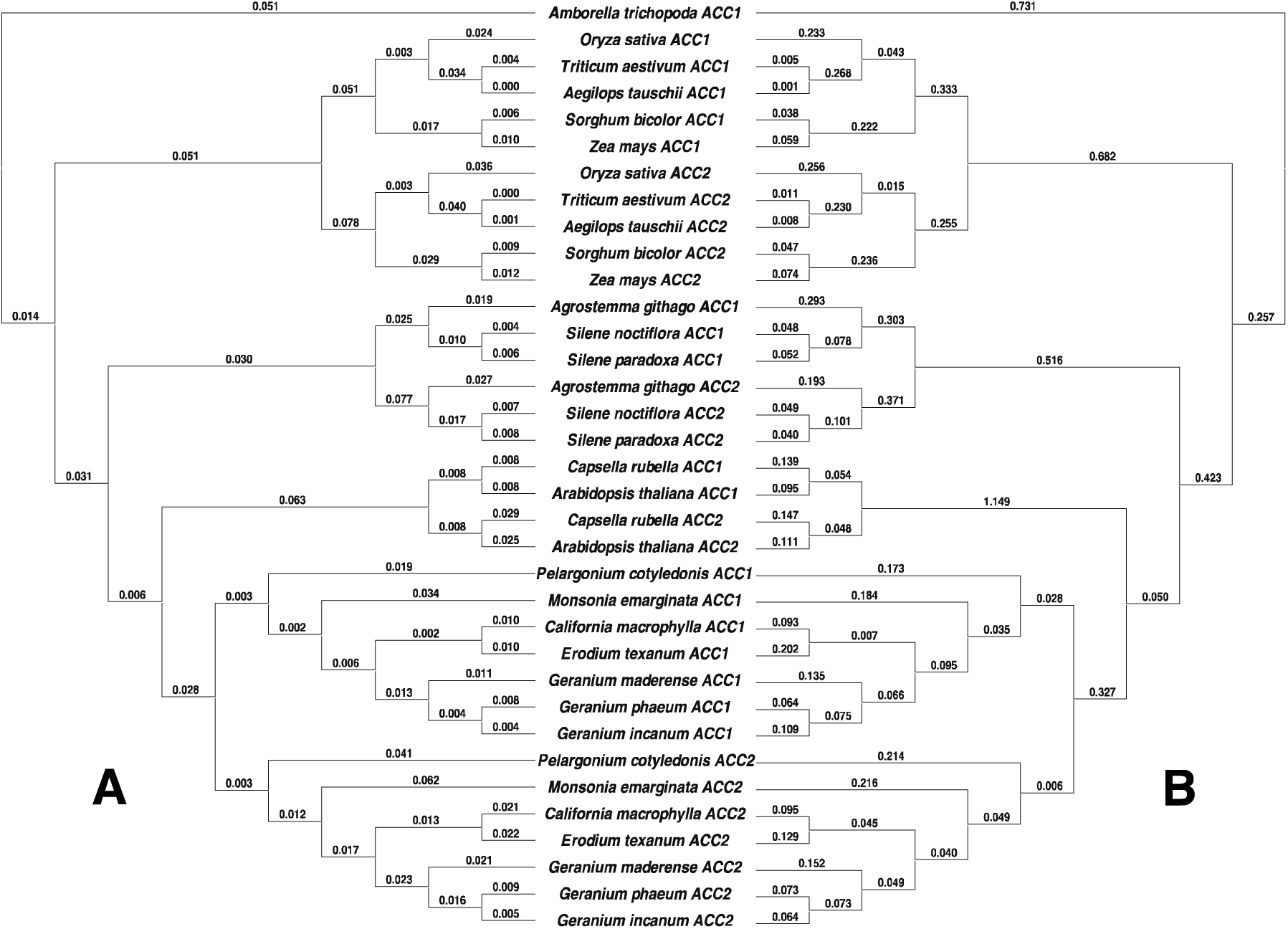
*ACC* tree. A) Branch labels are *d_N_* values. B) Branch labels are *d_S_* values.

**Figure S3.**
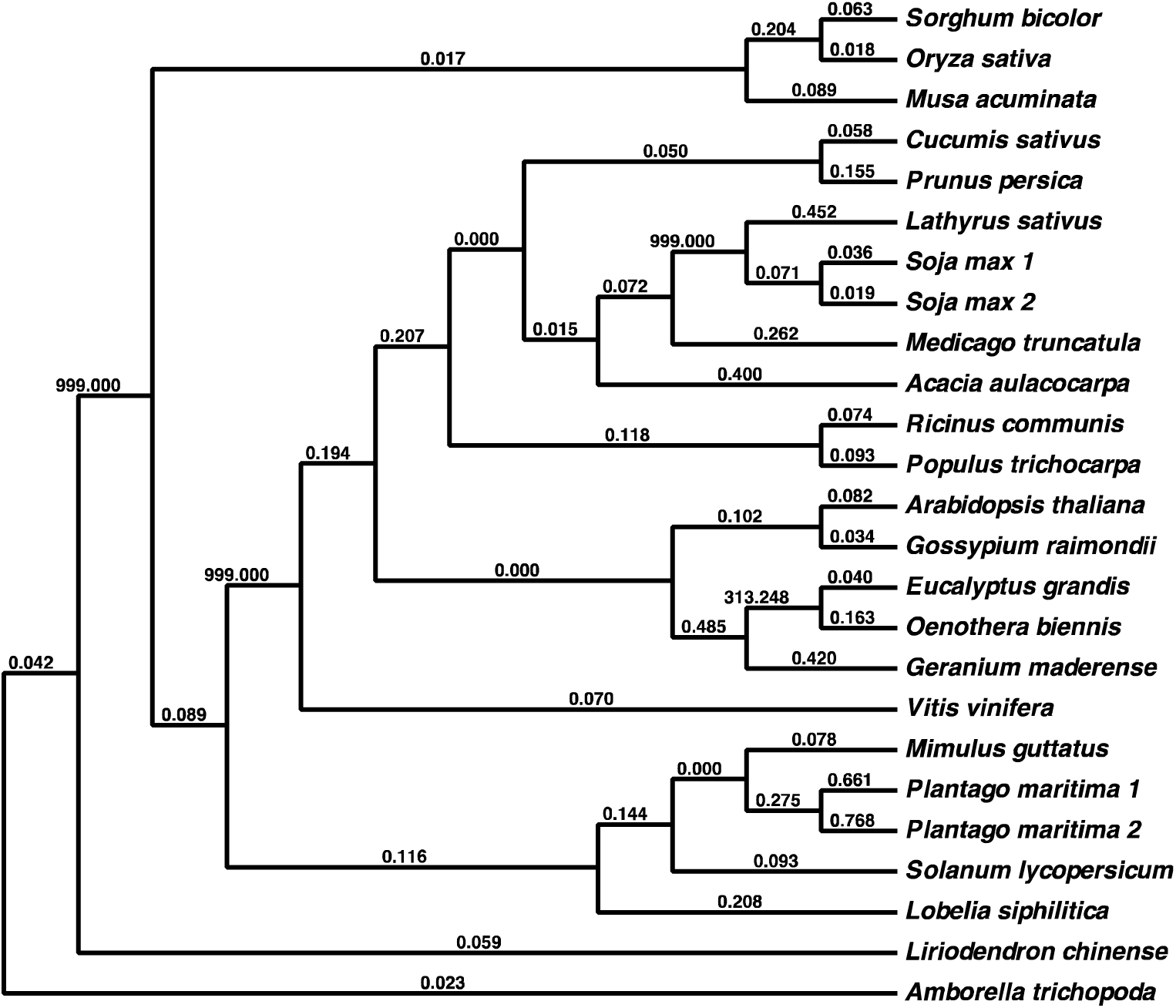
*CLPP3* tree. Branch labels are *d_N_*/*d_S_* values.

**Figure S4.**
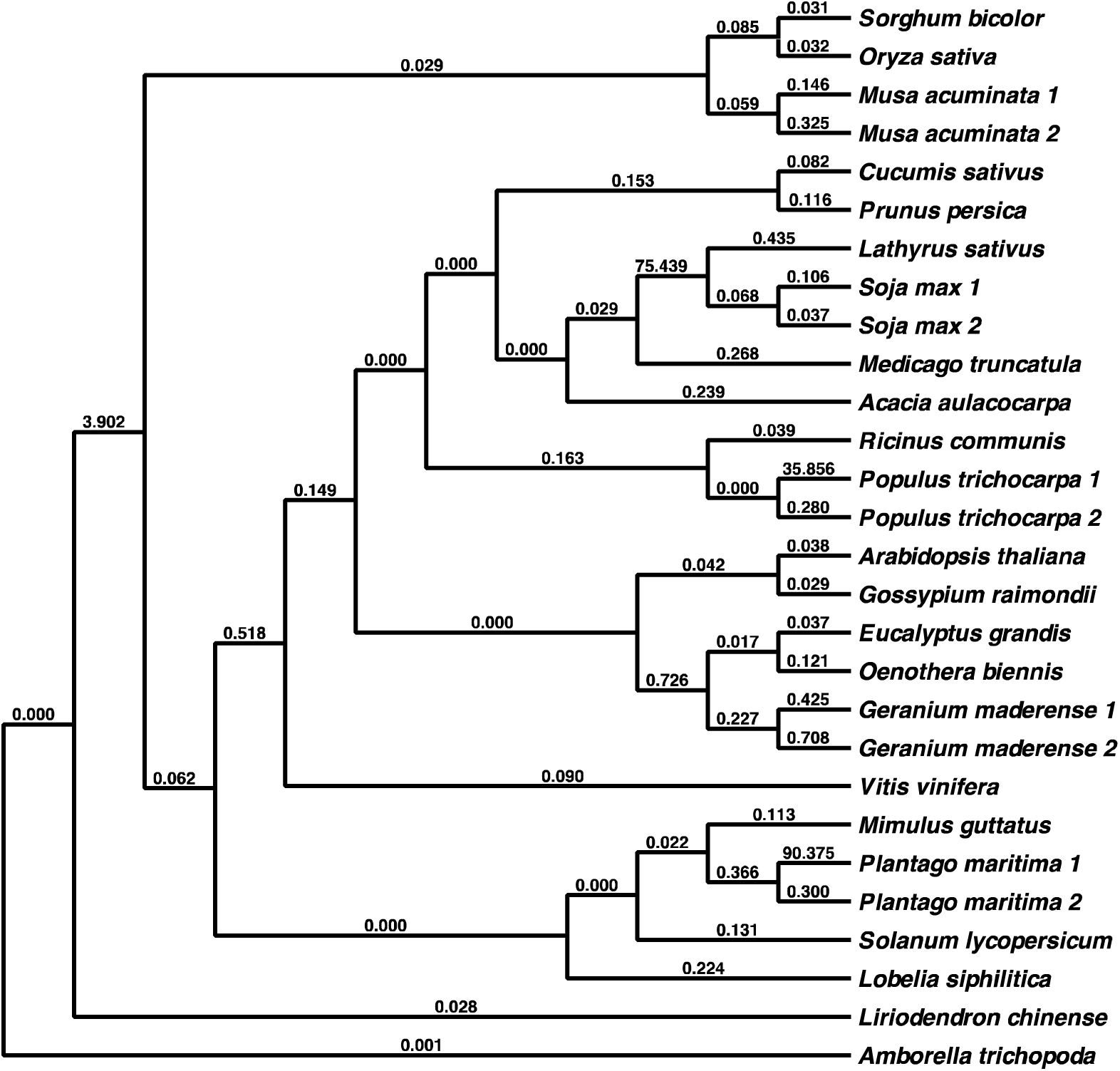
*CLPP4* tree. Branch labels are *d_N_*/*d_S_* values.

**Figure S5.**
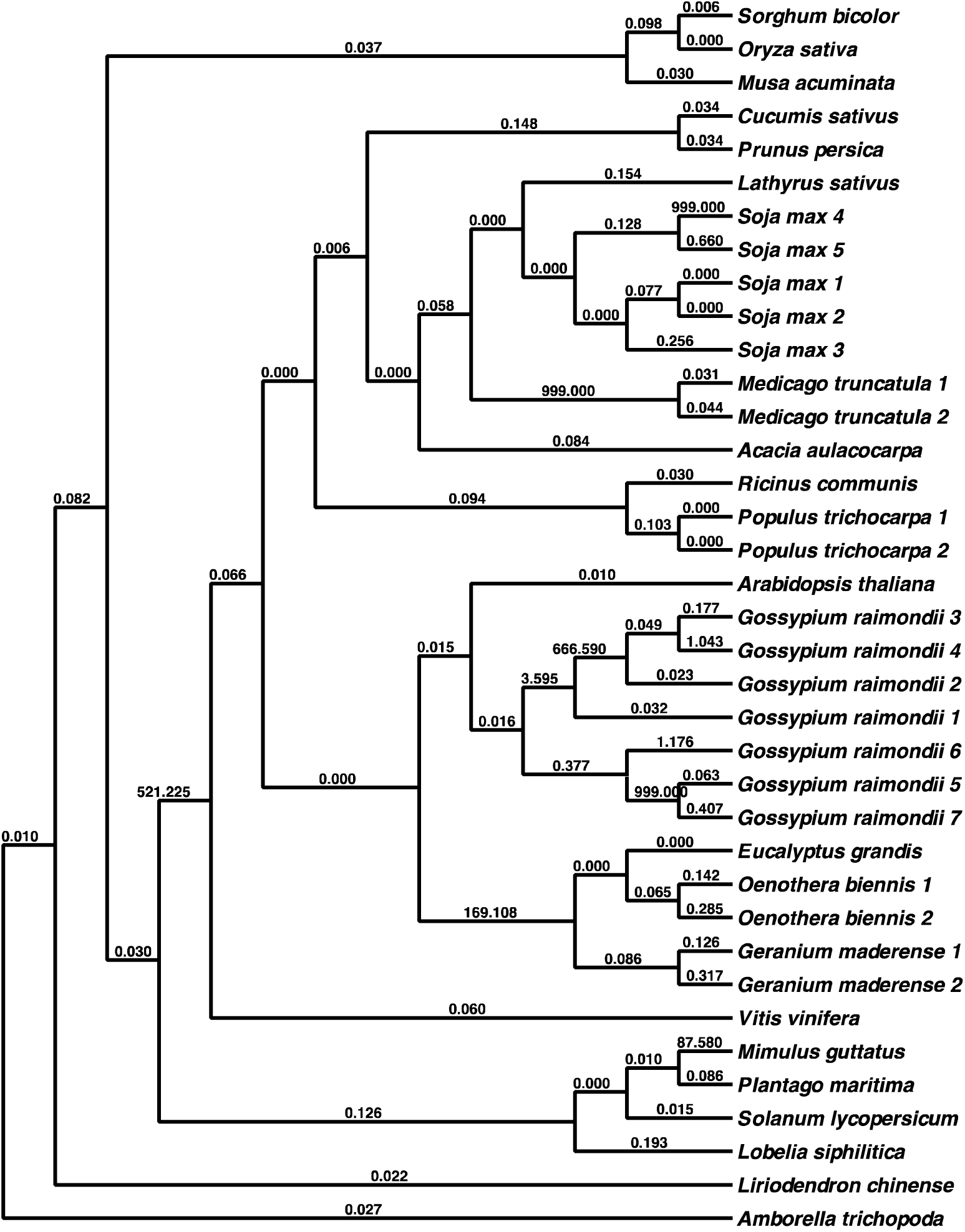
*CLPP5* tree. Branch labels are *d_N_*/*d_S_* values.

**Figure S6.**
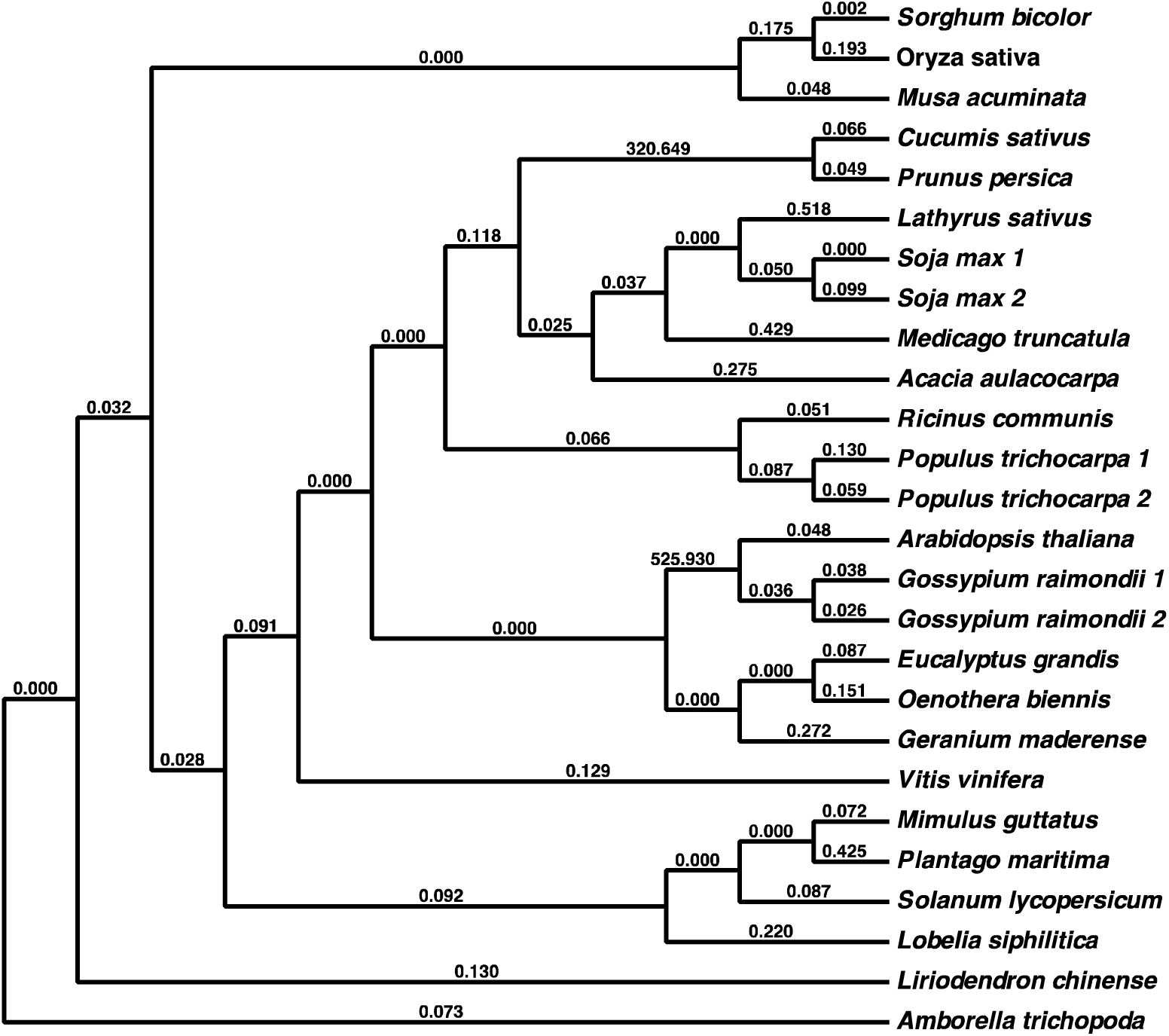
*CLPP6* tree. Branch labels are *d_N_*/*d_S_* values.

**Figure S7.**
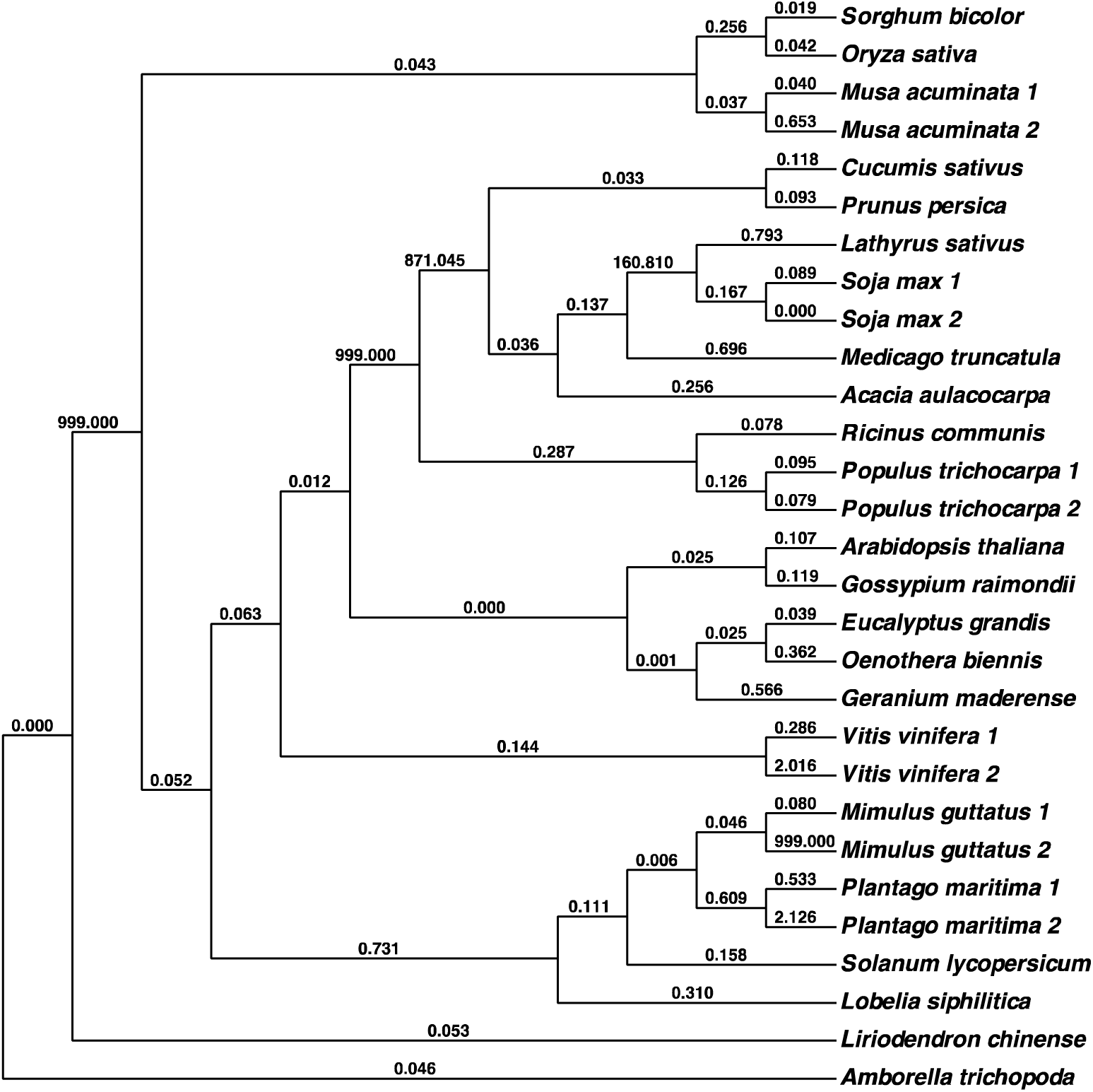
*CLPR1* tree. Branch labels are *d_N_*/*d_S_* values.

**Figure S8.**
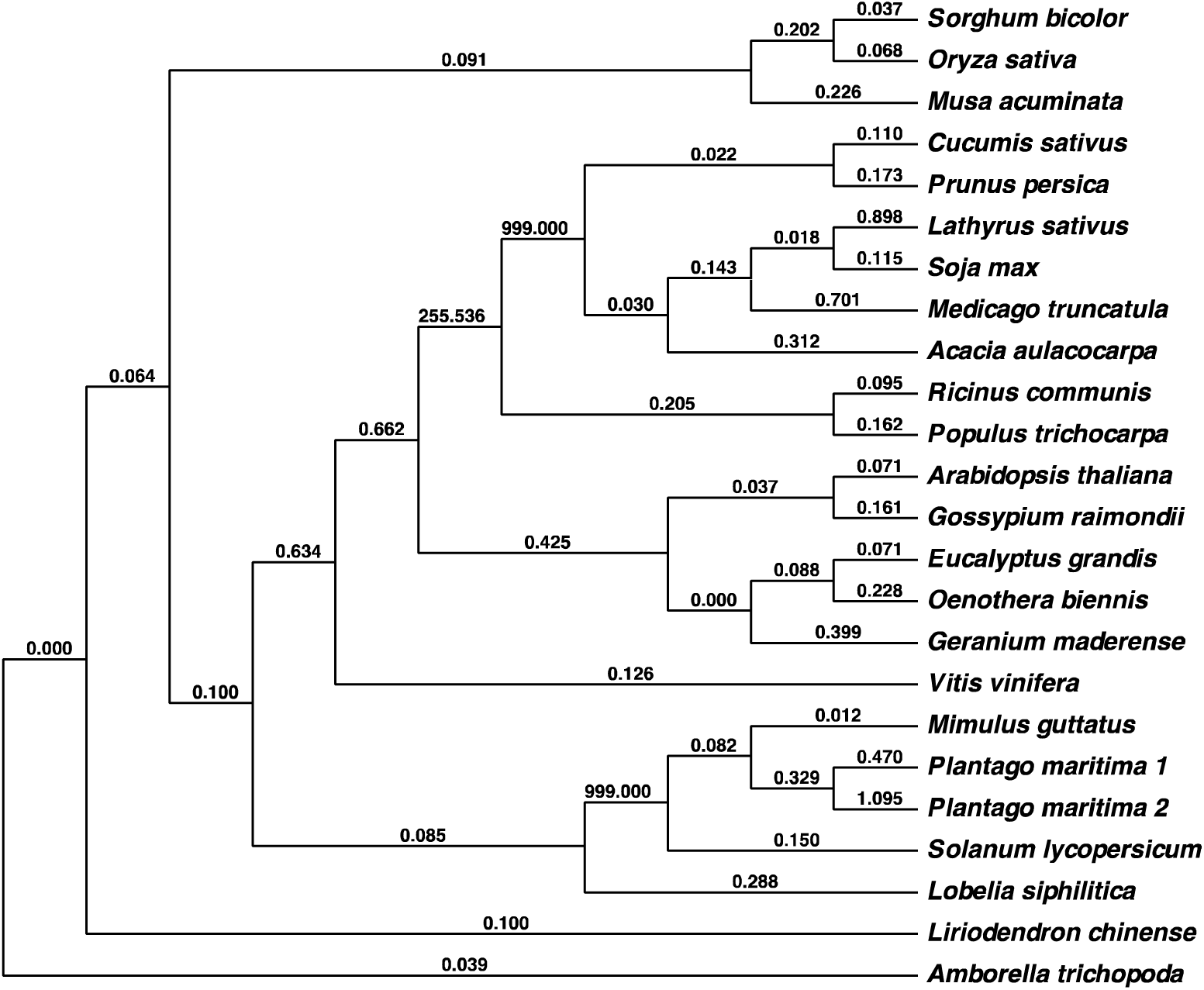
*CLPR2* tree. Branch labels are *d_N_*/*d_S_* values.

**Figure S9.**
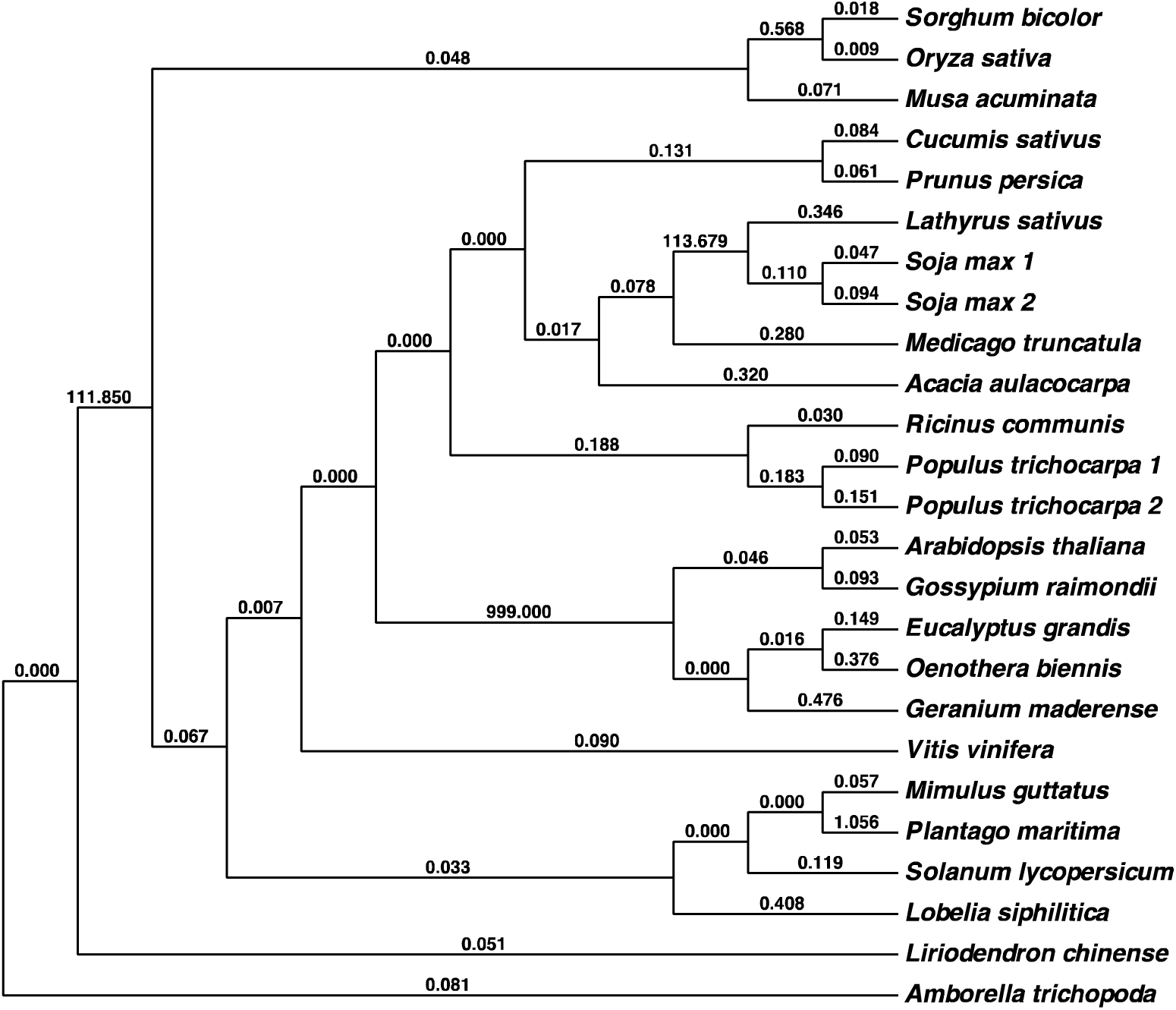
*CLPR3* tree. Branch labels are *d_N_*/*d_S_* values.

**Figure S10.**
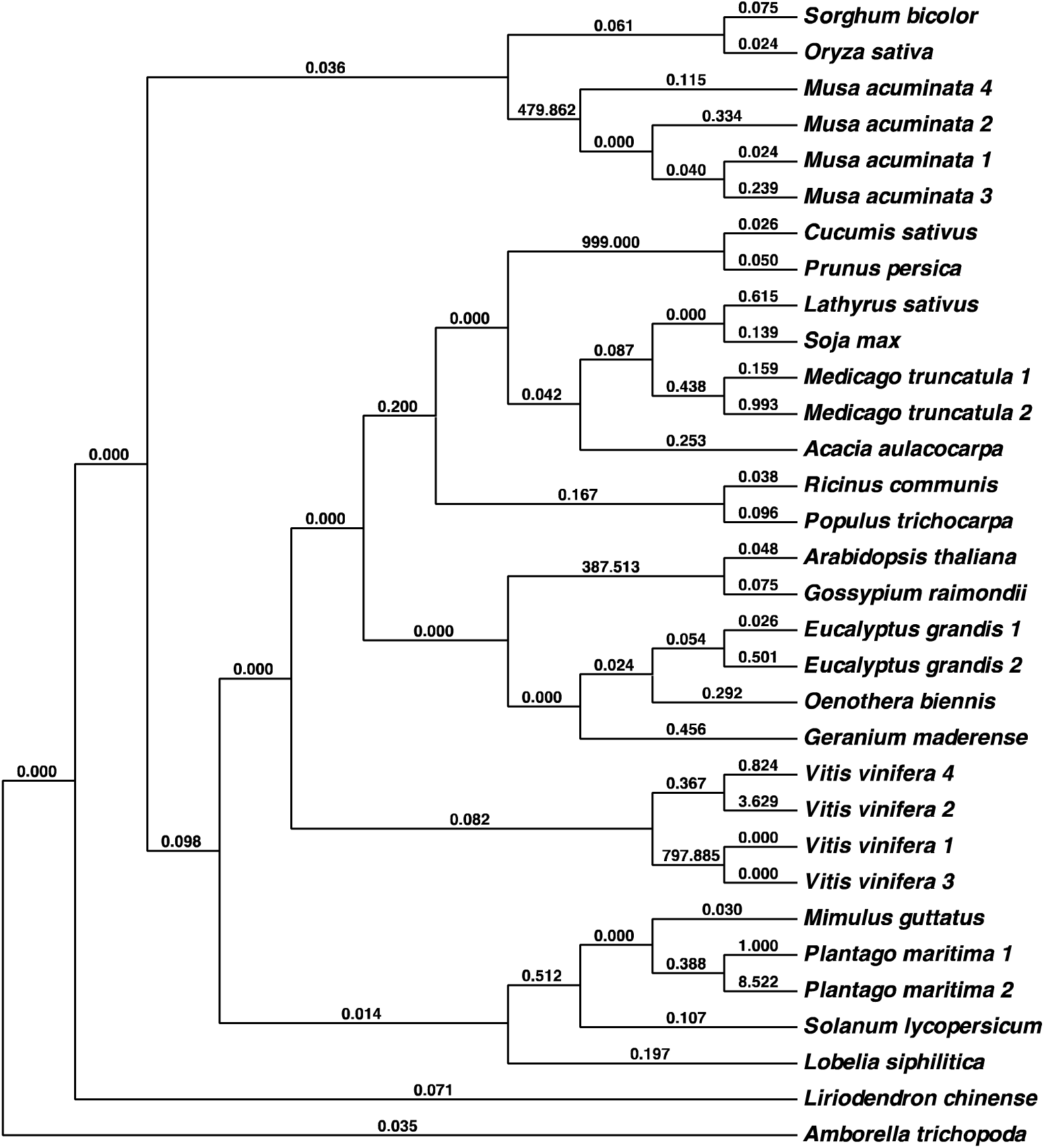
*CLPR4* tree. Branch labels are *d_N_*/*d_S_* values.

**Table S1:**
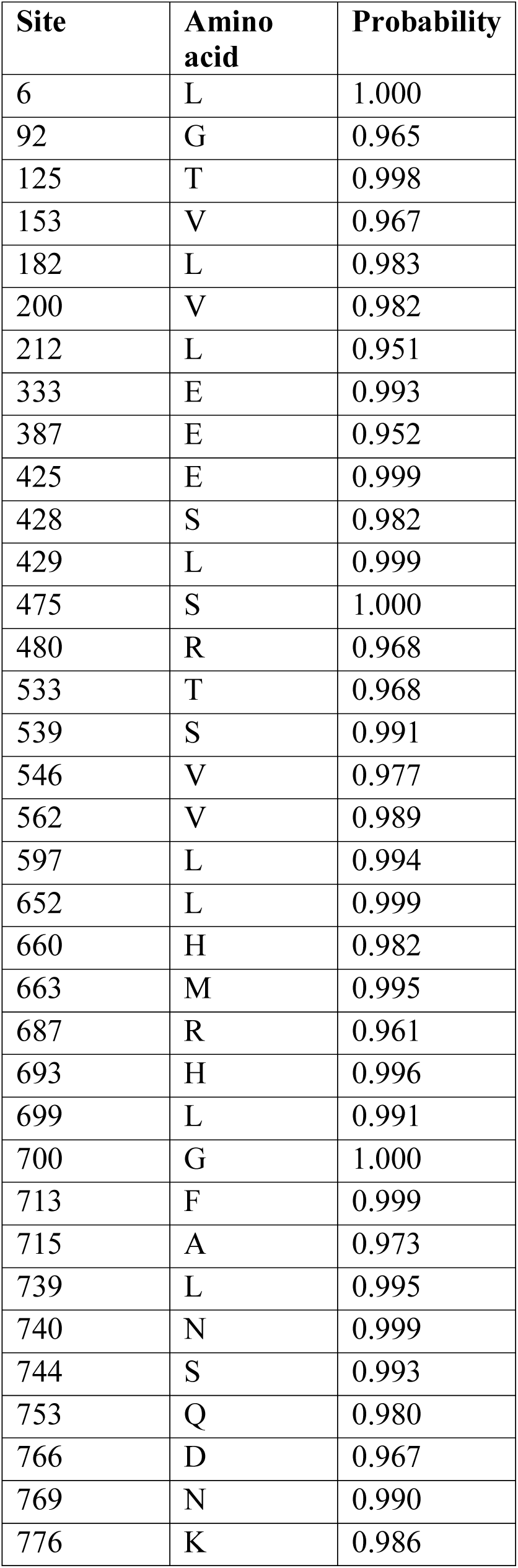
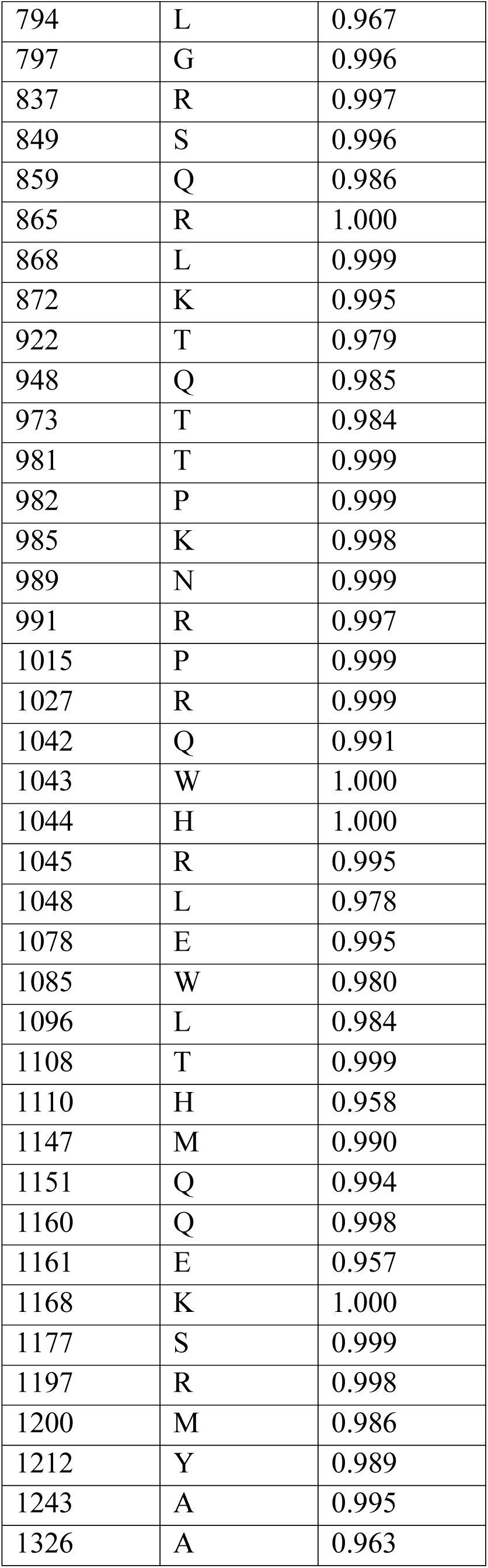
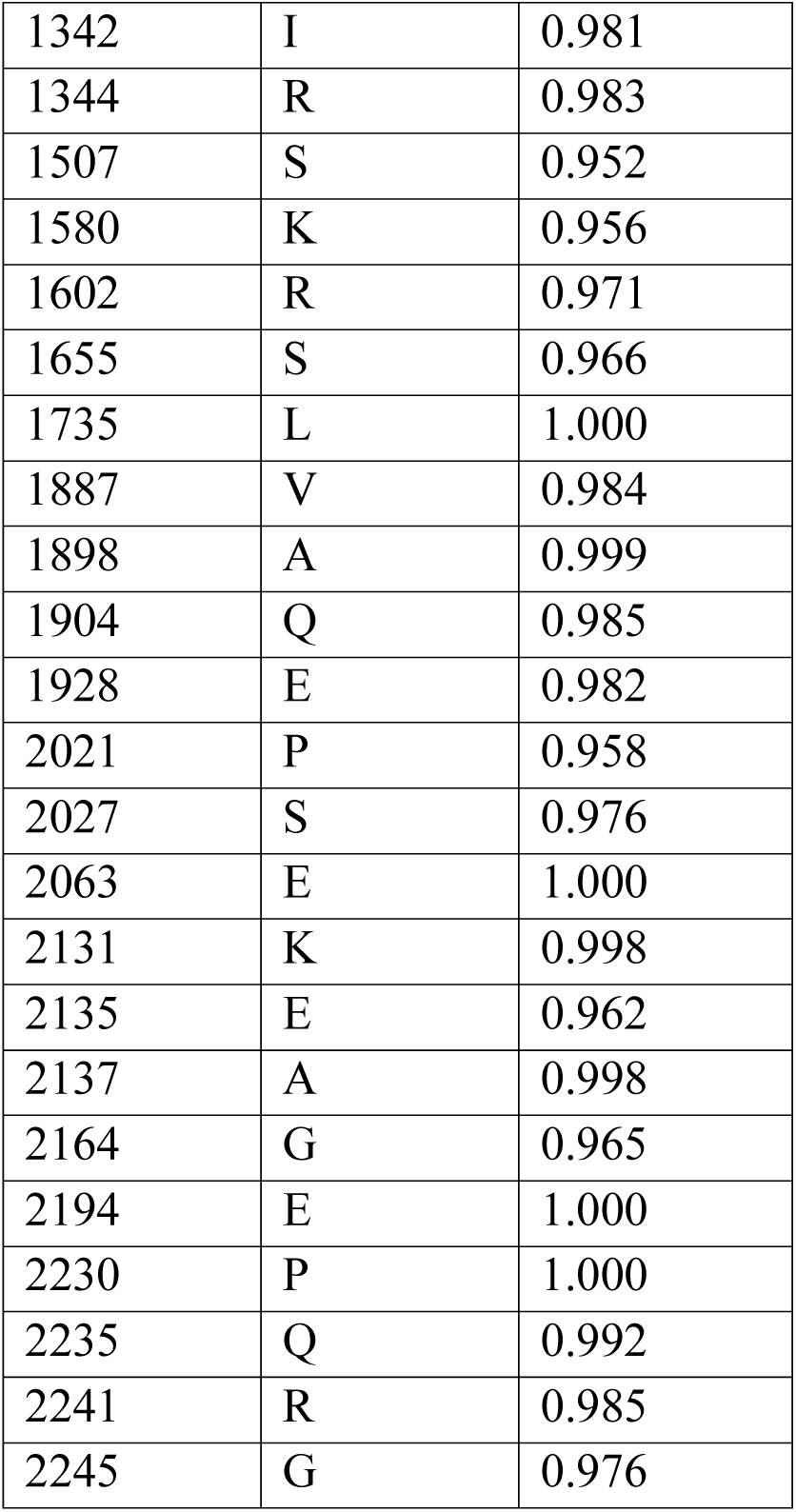
Sites inferred to be under positive selection in *ACC2* branches based on a branch-sites test in PAML using the trimmed alignment for *ACC*.

| Site | Amino acid | Probability |
| --- | --- | --- |
| 6 | L | 1.000 |
| 92 | G | 0.965 |
| 125 | T | 0.998 |
| 153 | V | 0.967 |
| 182 | L | 0.983 |
| 200 | V | 0.982 |
| 212 | L | 0.951 |
| 333 | E | 0.993 |
| 387 | E | 0.952 |
| 425 | E | 0.999 |
| 428 | S | 0.982 |
| 429 | L | 0.999 |
| 475 | S | 1.000 |
| 480 | R | 0.968 |
| 533 | T | 0.968 |
| 539 | S | 0.991 |
| 546 | V | 0.977 |
| 562 | V | 0.989 |
| 597 | L | 0.994 |
| 652 | L | 0.999 |
| 660 | H | 0.982 |
| 663 | M | 0.995 |
| 687 | R | 0.961 |
| 693 | H | 0.996 |
| 699 | L | 0.991 |
| 700 | G | 1.000 |
| 713 | F | 0.999 |
| 715 | A | 0.973 |
| 739 | L | 0.995 |
| 740 | N | 0.999 |
| 744 | S | 0.993 |
| 753 | Q | 0.980 |
| 766 | D | 0.967 |
| 769 | N | 0.990 |
| 776 | K | 0.986 |
| 794 | L | 0.967 |
| 797 | G | 0.996 |
| 837 | R | 0.997 |
| 849 | S | 0.996 |
| 859 | Q | 0.986 |
| 865 | R | 1.000 |
| 868 | L | 0.999 |
| 872 | K | 0.995 |
| 922 | T | 0.979 |
| 948 | Q | 0.985 |
| 973 | T | 0.984 |
| 981 | T | 0.999 |
| 982 | P | 0.999 |
| 985 | K | 0.998 |
| 989 | N | 0.999 |
| 991 | R | 0.997 |
| 1015 | P | 0.999 |
| 1027 | R | 0.999 |
| 1042 | Q | 0.991 |
| 1043 | W | 1.000 |
| 1044 | H | 1.000 |
| 1045 | R | 0.995 |
| 1048 | L | 0.978 |
| 1078 | E | 0.995 |
| 1085 | W | 0.980 |
| 1096 | L | 0.984 |
| 1108 | T | 0.999 |
| 1110 | H | 0.958 |
| 1147 | M | 0.990 |
| 1151 | Q | 0.994 |
| 1160 | Q | 0.998 |
| 1161 | E | 0.957 |
| 1168 | K | 1.000 |
| 1177 | S | 0.999 |
| 1197 | R | 0.998 |
| 1200 | M | 0.986 |
| 1212 | Y | 0.989 |
| 1243 | A | 0.995 |
| 1326 | A | 0.963 |
| 1342 | I | 0.981 |
| 1344 | R | 0.983 |
| 1507 | S | 0.952 |
| 1580 | K | 0.956 |
| 1602 | R | 0.971 |
| 1655 | S | 0.966 |
| 1735 | L | 1.000 |
| 1887 | V | 0.984 |
| 1898 | A | 0.999 |
| 1904 | Q | 0.985 |
| 1928 | E | 0.982 |
| 2021 | P | 0.958 |
| 2027 | S | 0.976 |
| 2063 | E | 1.000 |
| 2131 | K | 0.998 |
| 2135 | E | 0.962 |
| 2137 | A | 0.998 |
| 2164 | G | 0.965 |
| 2194 | E | 1.000 |
| 2230 | P | 1.000 |
| 2235 | Q | 0.992 |
| 2241 | R | 0.985 |
| 2245 | G | 0.976 |

**Table S2:** Loss of catalytic sites and truncation of nuclear-encoded plastid ClpP core subunits

| Species | Protein | Serine | Histidine | Aspartate | Length |
| --- | --- | --- | --- | --- | --- |
| *Eucalyptus grandis* | ClpP3 |  | Replaced with R |  |  |
| *Musa acuminata* | ClpP3 |  | Replaced with R |  |  |
| *Plantago maritima* 1 | ClpP3 | Replaced with Y | Replaced with R | Replaced with N |  |
| *Plantago maritima* 2 | ClpP3 | Replaced with Y | Replaced with R | Replaced with N |  |
| *Lathyrus sativus* | ClpP4 |  | Replaced with T |  |  |
| *Medicago truncatula* | ClpP4 |  | Replaced with A |  |  |
| *Musa acuminata* 2 | ClpP4 | Gap | Replaced with R | Gap | Truncated |
| *Plantago maritima* 2 | ClpP4 |  |  | Gap |  |
| *Populus trichocarpa* 2 | ClpP4 | Replaced with M | Gap | Gap | Truncated |
| *Gossypium raimondii* 3 | ClpP5 | Gap |  |  |  |
| *Gossypium raimondii* 4 | ClpP5 |  |  |  | Truncated |
| *Gossypium raimondii* 5 | ClpP5 | Replaced with N |  |  | Truncated |
| *Gossypium Raimondii* 6 | ClpP5 | Replaced with N |  | Replaced with N | Truncated |
| *Gossypium raimondii* 7 | ClpP5 | Replaced with N |  | Replaced with F |  |
| *Plantago maritima* | ClpP5 | Gap | Gap | Gap | Truncated |
| *Lathyrus sativus* | ClpP6 |  | Replaced with G |  |  |
| *Lobelia siphilitica* | ClpP6 | Replaced with N |  |  |  |
| *Medicago truncatula* | ClpP6 |  | Replaced with N |  |  |
| *Plantago maritima* | ClpP6 | Replaced with G | Replaced with E | Replaced with F |  |
| *Populus trichocarpa* 2 | ClpP6 |  |  |  | Truncated |

\*Empty cell indicates presence of catalytic site or full length.

**Table S3:**
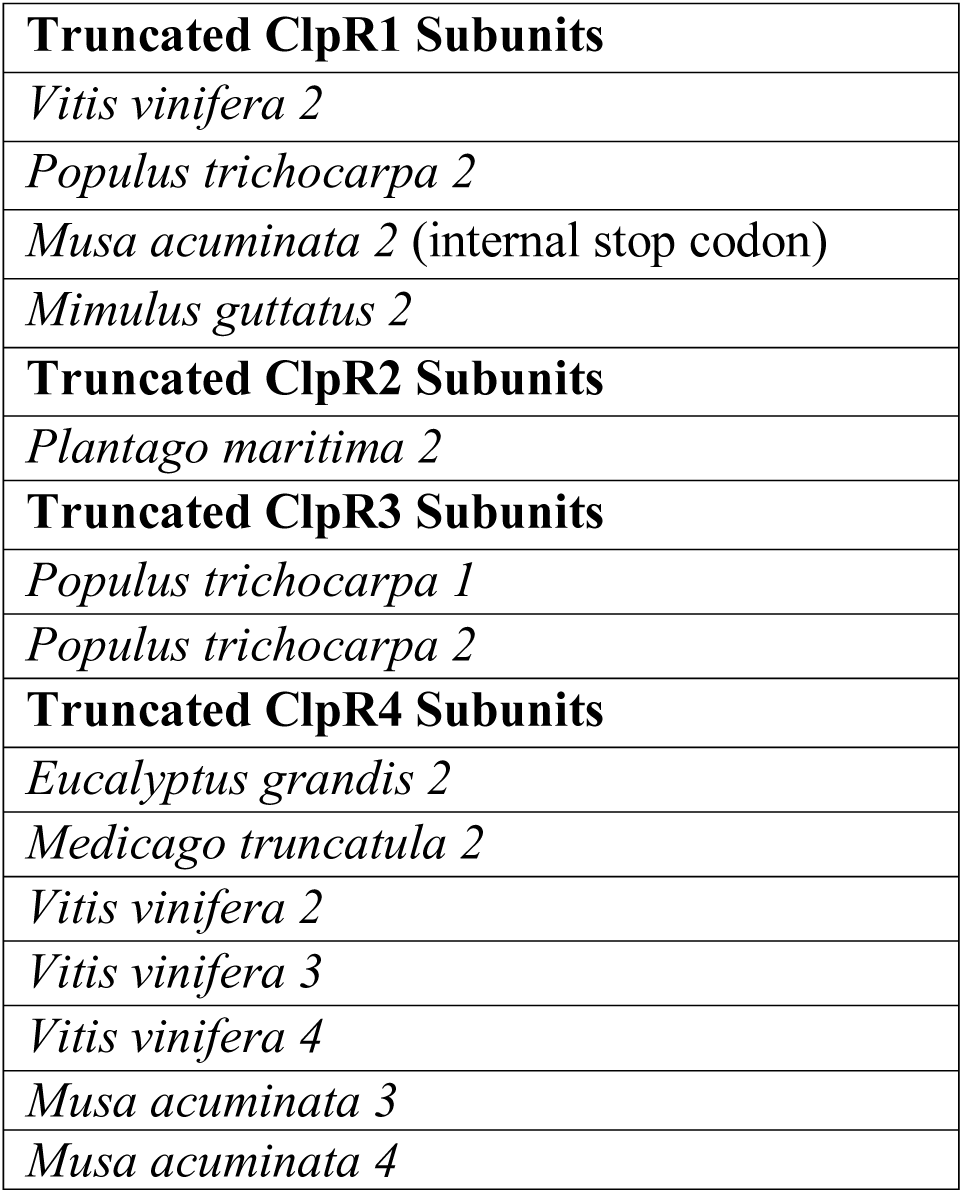
Truncation of nuclear-encoded plastid ClpR core subunits

